# Antimicrobial cetylpyridinium chloride disrupts the mitochondrial electron transport chain as potently as cyanide, via cardiolipin interference at cytochrome C

**DOI:** 10.64898/2026.08.18.745331

**Authors:** Emily L. Ledue, Noah E. Adelman, Madeleine K. Lorenger, Dylan J. Wagner, Sophie K. Trafton, Esther Biro, Emma R. Morrison, Quinn W. D’Alessio, John E. Burnell, Julie A. Gosse

## Abstract

People are widely exposed to the antimicrobial cetylpyridinium chloride (CPC) via consumer products, but CPC is a mitochondrial toxicant with potency comparable to that of canonical mitotoxicants. CPC is largely unregulated despite growing usage, bioavailability, and ability to cross the blood-brain barrier. Previously, we showed, in several cell types at non-cytotoxic and exposure-relevant doses, CPC inhibits ATP and OCR, endpoints of the electron transport chain (ETC). Mitochondrial toxicity is linked to multiple diseases (*e.g.*, diabetes, Parkinson’s, myalgic encephalomyelitis), but CPC has not been studied epidemiologically, and little mechanistic information is available. To determine why OCR and ATP are hampered by CPC, we hypothesized that CPC inhibits individual ETC components, cardiolipin, or TCA enzymes. Here, we show that, in primary human skin cells, an immune mast cell model, and isolated mitochondria, CPC apparently inhibits multiple ETC Complexes. Detailed investigation pinpointed the mechanism to the distal end of ETC: Complex III-cytochrome C-Complex IV. Using multiple approaches, we show that CPC does not directly inhibit any of the Complexes (not even Complex I as earlier reported), nor TCA enzymes, nor coenzyme Q. Yet, we found that CPC exhibits mitotoxicity as potent as cyanide. Anionic lipid cardiolipin attracts cytochrome C to the inner mitochondrial membrane so that it may shuttle electrons from Complex III to IV. Despite not altering levels of cardiolipin, CPC hinders cytochrome C by electrostatically interfering with cardiolipin. To aid epidemiology, risk analysis, and predictive toxicology, we have determined the precise biochemical mechanism of action of this ubiquitous compound.

## Introduction

People are widely exposed to an antibacterial agent that is as potent as canonical mitochondrial toxicants: the quaternary ammonium compound (quat) cetylpyridinium chloride (CPC) (Weller et al., 2024). CPC is found, increasingly (Fortune Business Insights, 2020), at millimolar (Franz, 2023) levels in numerous personal care (shampoos, mouthwash, lozenges, cosmetics) cleaning, pharmaceutical, and food such as in poultry processing (European Commission. Directorate General for Health and Food Safety., 2015; *HHS. Department Of Health And Human Services: Food And Drug Administration: Rules and Regulations: Secondary Direct Food Additives Permitted in Food for Human Consumption. [FR DOC # 04-7399] Final Rule 69, 17297–17298 (2004).*, 2020; Mao et al., 2020)(Pubchem;FDA 2020). As an example of its increasing use, CPC was recently employed in a newly developed urinary tract infection therapy, VesiX (Sawant et al., 2024). Exposure-relevant, non-cytotoxic CPC doses potently perturb mitochondrial morphology (as imaged with live-cell super-resolution microscopy), (Chávez & Bravo, 1982b; Liu et al., 2023; Saladino et al., 1971) as well as ATP,(Datta, He, et al., 2017a; Saladino et al., 1971; Weller et al., 2024) by inhibiting oxygen consumption rate (OCR)(Chávez & Bravo, 1982b; Datta, He, et al., 2017a; Weller et al., 2024), but the underlying biochemistry is unknown. CPC exerts its mitochondrial toxicity at ∼1000-fold lower doses than those currently encountered in products intended for human use. Abnormal mitochondrial function and morphologies are linked to numerous diseases(Youle & Van Der Bliek, 2012), such as diabetes(Jheng et al., 2012), memory loss(Hara et al., 2014), Parkinson’s, (Bhandari et al., 2014; Cui et al., 2010; Pozo Devoto & Falzone, 2017) COPD(Ryter et al., 2018), myalgic encephalomyelitis(Arron et al., 2024; Tomas et al., 2017).

CPC decreases OCR(Chávez & Bravo, 1982b; Saladino et al., 1971; Weller et al., 2024)--the mechanism of this suppression remains unknown. Previous reports indicated Complex I (NADH dehydrogenase) as CPC’s target in the electron transport chain (ETC)(Chávez & Bravo, 1982b; Datta, He, et al., 2017a; Saladino et al., 1971), but the mode of action remains uncertain. Previous studies used high CPC doses (100 μM, likely cytotoxic(Weller et al., 2024)) or cells with a Complex I mutation predisposing them to mitotoxicity, or relied on OCR endpoint, which necessarily integrates effects on all ETC components not just Complex I ((Baracca & Solaini, 2005; Datta, He, et al., 2017a; Kirches, 2011). That is to say, apparent inhibition of Complex I could be explained by chemical targeting of a downstream ETC component. ETC inhibitors are a known cause of mitochondrial fragmentation(Ahmad et al., 2013; Bulthuis et al., 2019; Giedt et al., 2012; Plecitá-Hlavatá et al., 2008); thus, determining how CPC dysregulates the ETC may also explain the structural defects. Here, we have systematically investigated CPC impacts on the complete ETC, to determine CPC’s mechanism of action.

A recent review called for research on this under-studied but large class of chemicals, quaternary ammonium compounds(Arnold & Blum, 2023), characterized by a permanently positively-charged head group and a lipophilic hydrocarbon tail (CPC structure in Fig. 10C). Among quats, CPC may be maximally toxic to eukaryotes, due to its 16 C chain length(Moss & Mayo-Bean, 2010; Zheng et al., 2021). For example, CPC is more potently mitotoxic than the better-studied benzalkonium chloride(Datta, Baudouin, et al., 2017a). Mechanistic information on CPC likely is applicable to other quats.

The use of quats has recently surged(Zheng et al., 2021) as a Covid treatment (Bañó-Polo et al., 2022; Koch-Heier et al., 2021). This is evidenced by a 331% post-pandemic increase in quats detected in wastewater treatment plants compared to before the 2019 SARS-CoV-2 outbreak(Mohapatra et al., 2023). Use of CPC as an antiviral against SARS-CoV-2 and influenza(Alemany et al., 2022; Alzahrani et al., 2023; Bezinelli et al., 2023; Bonn et al., 2023; Ferrer et al., 2021; Onozuka et al., 2024; Perussolo et al., 2023; Saud et al., 2022; Seneviratne et al., 2021; Tarragó-Gil et al., 2023) is an emerging exposure. Aggressive recent testing of CPC as an antiviral has indicated CPC destruction of lipid viral envelopes including SARS-CoV-2(Anderson et al., 2022; Bañó-Polo et al., 2022; Koch-Heier et al., 2021; Muñoz-Basagoiti et al., 2021; Popkin et al., 2017; Saud et al., 2022; Takeda et al., 2022), which is enriched in phosphatidylinositols(Saud et al., 2022), suggesting CPC interaction with these negatively-charged eukaryotic lipids. Since both SARS-CoV-2(Guarnieri et al., 2023) and CPC are toxic to mitochondria, benefits and risks of CPC use must be weighed. Through Toxic Substances Control Act loopholes, CPC and other quats have been exempted by the U.S. EPA from toxicity testing. However, CPC is recognized by the state of California as a Priority Chemical for biomonitoring. While the U.S. FDA has effectively banned CPC specifically from hand soaps and sanitizers (FDA, 2019), the chemical remains in widespread use in other products, noted above.

CPC works as an antibacterial by lysing microbes via detergent action when above its critical micelle concentration (CMC, ∼900 uM(Abezgauz et al., 2010; Mandal & Nair, 1991; Mukerjee & Mysels, 1971; Shi et al., 2011; Varade et al., 2005) doses. However, our studies use CPC doses ∼*100- to 10,000-fold* lower than the CMC, thus report effects that are *not* due to detergent lysis(Obeng et al., 2023, 2024; Raut, Weller, et al., 2022; Weller et al., 2024). For example, measures of plasma membrane integrity (lactate dehydrogenase release(Raut, Weller, et al., 2022) and plasma membrane potential(Obeng et al., 2023) are unaffected by the CPC exposure conditions used in this study. All utilized exposures (dose, time) were non-cytotoxic, including via trypan blue and ToxGlo assays(Obeng et al., 2023; Raut, Weller, et al., 2022; Weller et al., 2024).

Previous work from our research group showed that CPC potency is on par with the canonical mitotoxicant carbonyl cyanide 3-chlorophenyl hydrazone (CCCP) and much more potent than other well-described mitotoxicants, triclosan (Weatherly et al., 2016) and 2,4-dinitrophenol(Weller et al., 2024). We found CPC is a mitotoxicant in three cell types from different species and tissue types, including primary human keratinocytes, at exposure-relevant levels(Bonesvoll & Gjermo, 1978; Obeng et al., 2023). As a striking example, CCCP inhibits ATP production with an IC_50_ of ∼1.2 μM(Weatherly et al., 2016), indicating that CPC (IC_50_ ∼1.7 μM) is as mitotoxic as this canonical uncoupler. Banned in 1938(Grundlingh et al., 2011), 2,4-dinitrophenol (DNP)(Loomis & Lipmann, 1948) is ∼184-fold *less* potent than CPC.

Despite widespread use, there is a lack of exposure studies directly quantifying levels of CPC in humans. However, our estimates derived from data (including the European Union Scientific Committee on Consumer Safety and a mouse pharmacokinetics study(Pottel et al., 2020)) suggest that human blood levels from acute CPC exposure may reach ∼0.3 uM from normal usage of personal care products with up to ∼0.3 uM more from a meal of CPC-treated chicken (possibly totaling ∼0.6 uM)(Weller et al., 2024). Furthermore, the pharmacokinetics of excretion are unknown, so the possibility remains that repeated exposures may be bioaccumulative. Following typical mouthwash use, CPC persists in the oral cavity, being slowly released into human saliva, at high-micromolar CPC concentrations for many hours/days after brief product usage(Bonesvoll & Gjermo, 1978) CPC also has been found in rivers (0.15 uM;(Shrivas & Wu, 2007)) and wastewater (0.25 uM)(Zaman et al., 2025). The concentrations used in this study range from 10 nM to 20 uM, so are directly comparable to real human and ecological exposure levels. Similar quats have been found in people (80% of humans tested in one study) including in breast milk(Zheng et al., 2022), and the quats reduced OCR in exposed humans(Hrubec et al., 2021)–but CPC was not tested in those studies. In searching (PubMed, 2026), we have found no other direct assessments of CPC body burden, apart from one study in which CPC was detected in the blood and urine of 1 out of 9 humans studied, but concentrations were not reported (EWG Human Toxome Project (2003).

Mounting evidence suggests CPC breaches the blood-brain barrier(Cohn et al., 2024) and is bioavailable(Klaassen & Casarett, 2019; Van Leeuwen et al., 2015; Voutchkova et al., 2010). Specifically, CPC has been noted to kill developing oligodendrocytes (IC_50_ = 68.5 nM)(Cohn et al., 2024). Decreased heart rate and swim velocity were observed in zebrafish exposed to 0.2 uM CPC in water(Qiu et al., 2022). CPC affects both tadpole survival rate (IC_50_= 2 uM) and embryonic development (1.5 uM; (Park et al., 2016)). At 0.18 uM, CPC induces oxidative stress in the sewage worm *Tubifex tubifex(Bhattacharya et al., 2021)*. In addition to these effects, CPC impairs specific immune cell signaling events in mast cells(Obeng et al., 2024).

Models used in this study reside in humans from the top to bottom of the oral mucosa/skin/digestive tract (primary human skin cells, mast cells, mitochondria). Mast cells (MC) are involved in many diseases and physiological processes(Galli et al., 2005; Metcalfe et al., 1997; Silver & Curley, 2013), and are ubiquitous, including at environmental interfaces like skin and oral/GI mucosa(Kuby, 1997), so may be exposed to CPC. The RBL-2H3 MC model is a widely accepted line that behaves like mature human MCs(Abramson & Pecht, 2007; Metcalfe et al., 1997; Metzger, 1982; Seldin et al., 1985) and responds to environmental stimuli similarly to primary MCs(Alsaleh et al., 2016; Thrasher et al., 2013). MC signaling(Obeng et al., 2024; Raut, Waters, et al., 2022a) and mitochondrial function(Weller et al., 2024) are inhibited by CPC. Primary human keratinocytes are a proxy for barrier cells, the last layer of protection between humans and our environment; these cells’ sensitive ATP and OCR responses to CPC were previously reported(Weller et al., 2024). Isolated mitochondria were also used to detail mechanistic processes. Regardless of tissue/eukaryotic species, the findings were consistent in this study and previously(Weller et al., 2024), likely due to the highly conserved nature of the mitochondrial ETC.

Electrons enter the ETC via two main donors: nicotinamide adenine dinucleotide (NADH) or flavin adenine dinucleotide (FADH_2_), from the tricarboxylic acid (TCA) and other sources, like -glycerol-phosphate dehydrogenase. Respectively, NADH and FADH_2_ feed electrons into Complex I and Complex II, both of which pass electrons next to coenzyme q (CoQ), from which they move on to Complex III, cytochrome c (cyt C), and finally to Complex IV, where electrons reach their terminal acceptor, O_2_.

The function of all ETC Complexes depends on the specific mitochondrial lipid cardiolipin (CL), a critical negatively charged phospholipid comprising up to 20% of all lipid content of the inner mitochondrial membrane (IMM)(De Kroon et al., 1997; Gebert et al., 2009). CL is composed of a glycerol backbone connecting to four acyl chains (Fig. 10C). CL aids in energy production by stabilizing the ETC complexes by binding each individual complex at specific sites(Pfeiffer et al., 2003). CL also binds cyt C(Jussupow et al., 2019; Musatov & Robinson, 2014; Schlame et al., 2000; Schwall et al., 2012), electrostatically pulling it from the aqueous intermembrane space down to the IMM surface through interactions between the anionic phosphate groups of CL and cationic amino acids on cyt C(Osheroff et al., 1983; Rytömaa et al., 1992; Rytömaa & Kinnunen, 1994; Vik et al., 1981). Thus, CL temporarily anchors cyt c to the IMM, facilitating the transfer of electrons from Complex III to IV(Hanske et al., 2012; Kawai et al., 2005; Sorice et al., 2009).

In this study, we aimed to uncover the biochemical mechanism by which CPC disrupts mitochondrial OCR. Here, we examined CPC effects on the members of the mitochondrial respiratory network, including the key lipid cardiolipin. We demonstrate that CPC, as potently as cyanide, impairs the ETC and does so not via Complex I but by targeting cardiolipin.

## Materials and Methods

Safety precautions, per each chemical’s safety data sheet, were used for each chemical and assay.

## Cetylpyridinium chloride monohydrate preparation

Cetylpyridinium chloride (CPC; CAS No. 6004-24-6, 99.7% purity; MP Biomedicals) was prepared in cell culture water (CCW; endotoxin-free, 0.1 uM sterile-filtered; VWR) as previously described(Raut, Weller, et al., 2022) via sonication, averting potential toxic effects that organic solvents may impose on cells. CPC was then diluted to the necessary experimental concentrations in the specified buffers used in each experimental condition. Due to its light sensitivity, CPC was utilized in experiments in low light conditions. CPC does not absorb UV–Vis light beyond ∼280 nm and thus will not interfere with the measurement of the probes used for Mitocheck assays (**Table S3**).

## Cell Culture

Both cell types were cultured as previously described: RBL-2H3 Rat Mast Cells(Hutchinson et al., 2011) and adult primary human epidermal keratinocytes(Weatherly et al., 2016).

## Biolog Electron Flow Assay

The Biolog Mitochondrial Assay Solution (MAS; 2X) is a minimal-nutrient media. MAS (at 2X) consists of: 260 mM Sucrose, 7.12 mM K_2_HPO_4_, 2.88 mM KH_2_PO_4_ (pH 7.4), 5 mM MgCl_2_, 1 mM EGTA (ethylene glycol-bis(2-aminoethylether)-N,N,N′,N’-tetra-acetic acid), 0.2% fatty acid free bovine serum albumin (GoldBio)(Lei & Bochner, 2021). For cell preparation, MAS was diluted to 1X in CCW. Saponin (CAS No. 8047-15-2, 8-25% sapogenin; Sigma Aldrich) was dissolved in CCW as a 24X solution. To permeabilize the plasma membrane for substrate entry without compromising mitochondrial membrane integrity and function, cells (similar to those used in this study) were treated with saponin at 30 ug/mL(Clerc & Polster, 2012; Coden et al., 2020; Lei & Bochner, 2021; Miyamoto et al., 2008; Salabei et al., 2014; Schulz, 1990). With this saponin dose focus is trained upon mitochondrial metabolism because 1.) TCA intermediates readily enter the cell and the integrity of the mitochondrial membrane remains intact and 2.) cytosolic metabolic substrates like glucose are not metabolized due to sufficient plasma membrane/cytosol disruption(Lei & Bochner, 2021).

*Substrate Reconstitution:* Biolog Mitoplate^TM^-S1 96-well plate well bottoms are coated with 31 substrate types (each at 5 mM in the final assay solution upon full dissolution), along with one no-substrate control well, all in triplicate. The substrates (and no-substrate control) were reconstituted by agitating the plate in activity buffer. To do so, activity buffer (2X in each component) was prepared as a master mix from concentrated solutions of the following (modified for each specified condition) in order to reach manufacturer’s suggested concentrations: CPC (or CCW), saponin, MAS and Redox Dye MC. Aliquots of this 2X activity buffer were dispensed at 30 uL/well into the Mitoplate^TM^-S1, which was incubated for 2 h at 37 °C/5% CO_2_. For robust substrate dissolution, the plate was agitated manually every 30 minutes during the 2 h incubation. After the 2 h, cells were added to each well as noted below.

*Cell preparation for Biolog assay:* Confluent RBL-2H3 cells from a T25 flask were trypsinized (0.05% trypsin with 0.53 mM EDTA, Gibco) at 37°C/5% CO_2_ for 5 min, quenched in RBL media, then centrifuged (TC Eppendorf 543 for 8 min at 300xg). The cell pellet (containing about 8 million cells) from the whole flask was then reconstituted into 3 mL of 1X MAS. Next, cells were strained through a 70-micron cell strainer (VWR), then counted. Strained cells were diluted into additional 1X MAS to a concentration of 1.33 x 10^6^ cells/mL and plated, 30 uL/well (40,000 cells/well), bringing the total final volume to 60 uL (including the 2X activity buffer) in each well.

Confluent primary human keratinocytes from a T25 flask were rinsed with phosphate buffered saline (PBS; Lifeline) and trypsinized 0.05% trypsin, 0.02% EDTA; Lifeline) at 37°C/5% CO_2_ for 5 min, quenched in Trypsin Neutralizing Solution (Lifeline), then centrifuged (TC Eppendorf 543 for 4.5 min at 150xg). The cell pellet (containing about 2 million cells) from the whole flask was then reconstituted into 2 mL of 1X MAS. Next, cells were diluted into additional 1X MAS to a concentration of 500,000 cells/mL and plated, 30 uL/well (15,000 cells/well), bringing the total final volume to 60 uL (including the 2X activity buffer) in each well.

*Electron Flow Detection:* The MC Redox Dye acts as a terminal electron acceptor, receiving electrons from cyt C after Complex III of cellular respiration along the mitochondrial inner membrane (Fig. 1). Dye reduction was then detected via A_590_ kinetically read (Synergy H1 plate reader, Agilent Biotek) over the course of ∼1-3 h at 37°C/5%CO_2_ and read into Gen5 2.0 software. Analyses were conducted to ensure that CPC does not affect the Biolog MC redox dye absorbance signal, such that it is possible to conclude that any effects due to CPC in Biolog electron flow data are due to physiological effects of CPC on the cells, rather than dye interference artifacts (Fig. S1).

**Figure 1.**
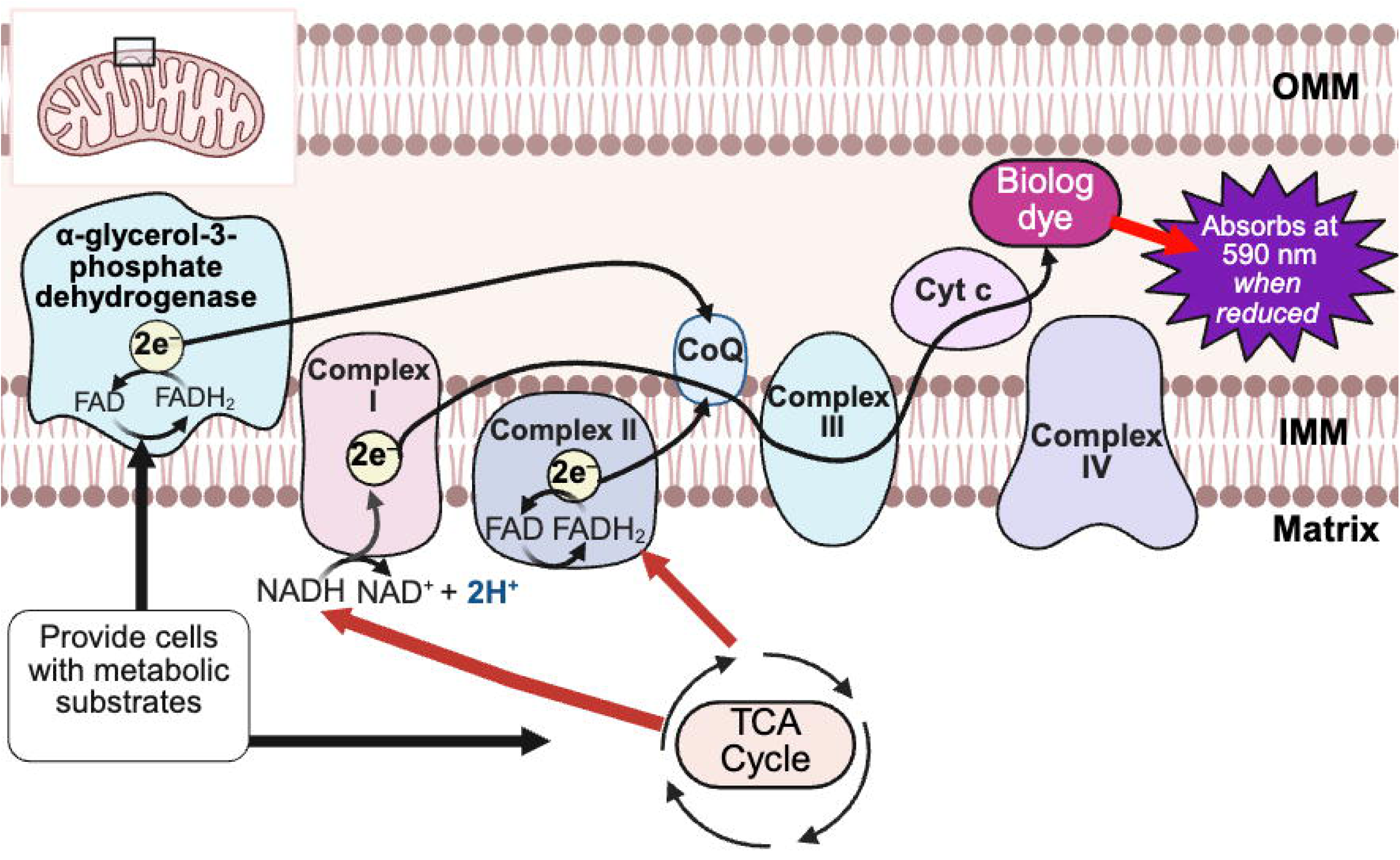
Biolog electron flow assay workflow. Permeabilized cells (RBL-2H3 or primary human keratinocytes) were fed either no substrate or 1 of 31 substrates. These substrates are metabolized inside mitochondria, such as via the tricarboxylic acid (TCA) cycle, into reducing power fed into the electron transport chain, eventually reducing Biolog MC redox dye instead of going on to complex IV (where oxygen is typically the terminal electron acceptor.) Successful electron flow is indicated by dye reduction. Dye reduction is observed as A_590_, read via plate reader. Rate of dye reduction from a particular metabolized substrate was compared in no-CPC control wells, versus 0.5 μM or 1.0 μM CPC wells. (OMM, outer mitochondrial membrane; IMM, inner mitochondrial membrane; Cyt C, cytochrome c; CoQ, coenzyme q). Created in Biorender.

*Data Analysis:* Absorbance controlling for abiotic background was then read at A_750_ for 5 min, time-averaged, then subtracted from each individual respective well’s A_590_ value. Next, the A_750_-subtracted A_590_ values of each no-substrate control well, within each CPC treatment group (0, 0.5, or 1 uM), were then deducted from each of the 31 substrate wells within each CPC treatment group, at every time point. See Table S1 for further analysis details regarding substrates selected for further analysis.

The linear region for each of the substrates which produced statistically significant electron flow (those in Table S1) in the control (0 uM CPC) group was determined via Microsoft Excel R^2^ function. A time-range corresponding to an R^2^ threshold (minimum R^2^= 0.87), individually determined for each substrate (Table S2), was used to calculate the electron flow slope (Excel slope function). This 0 uM CPC control slope was compared to the slope within the same time-range for the 0.5 uM CPC and 1.0 uM CPC groups. This was repeated for each experimental day (3-4 total experiments). The slope values for each experimental group were entered into Graphpad Prism for one-way ANOVA and Dunnett’s post-hoc test.

## Mitocheck Complex I Assay

CPC does not absorb UV–Vis light beyond ∼280 nm and thus will not interfere with the measurement of the probes used for Mitocheck assays (Table S3). *Rotenone Preparation for Complexes I, II, and III Investigations:* Rotenone (Cayman Chemical; CAS No. 83-79-4) is virtually insoluble in water (Pubchem), so rotenone was first dissolved in the organic solvent dimethyl sulfoxide (DMSO; Thermo Fisher Scientific, CAS No. 67-65-8) and then diluted with warmed Complex buffer (Complex I, II, and III buffers) to minimize the concentration of organic solvent. We found that rotenone must be prepared fresh every day for its efficacy. Immediately after dilution, rotenone solutions were vortexed and then added to samples. Vehicle controls were performed to ensure that the DMSO vehicle did not interfere with the assay (Fig. S3B). Rotenone is colorimetric; therefore, we needed to ensure there was no overlap between the wavelength of rotenone’s absorbance and the wavelengths assayed for the individual mitochondrial Complex analyses (Table S3). There is no significant overlap with rotenone’s absorbance (spectrum provided by Cayman Chemical, not shown) for the dyes in these assays.

*Mitocheck Complex I Data Collection:* MitoCheck® Complex I Activity assay kit (Cayman Chemical) measures electron flow from NADH through the full ETC (Fig. 4A). All materials used except for the treatment conditions were provided with the assay kit. Isolated bovine heart mitochondria were dissolved into in Complex I buffer with fatty acid free-bovine serum albumin as per manufacturer’s instructions, then were pre-exposed to each treatment group (CPC, positive control inhibitor rotenone, or vehicle control) for 60 min at 37 °C/5% CO_2_. Complex I activity was then measured in MitoCheck Complex I buffer with DMSO (0.05%) as required by the rotenone dissolution, in order to standardize the vehicle across all treatment types. Following the aforementioned 60 min pre-treatment, NADH and CoQ were added per manufacturer’s instructions. Complex I enzyme activity, correlating to the rate of NADH oxidation, was measured via kinetic A_340_ absorbance reading in the plate reader (Synergy H1 plate reader, Agilent Biotek) for 20 min at 12 s intervals and at 25 °C. Example background-subtracted results are shown in Fig. S3A.

**Figure 2.**
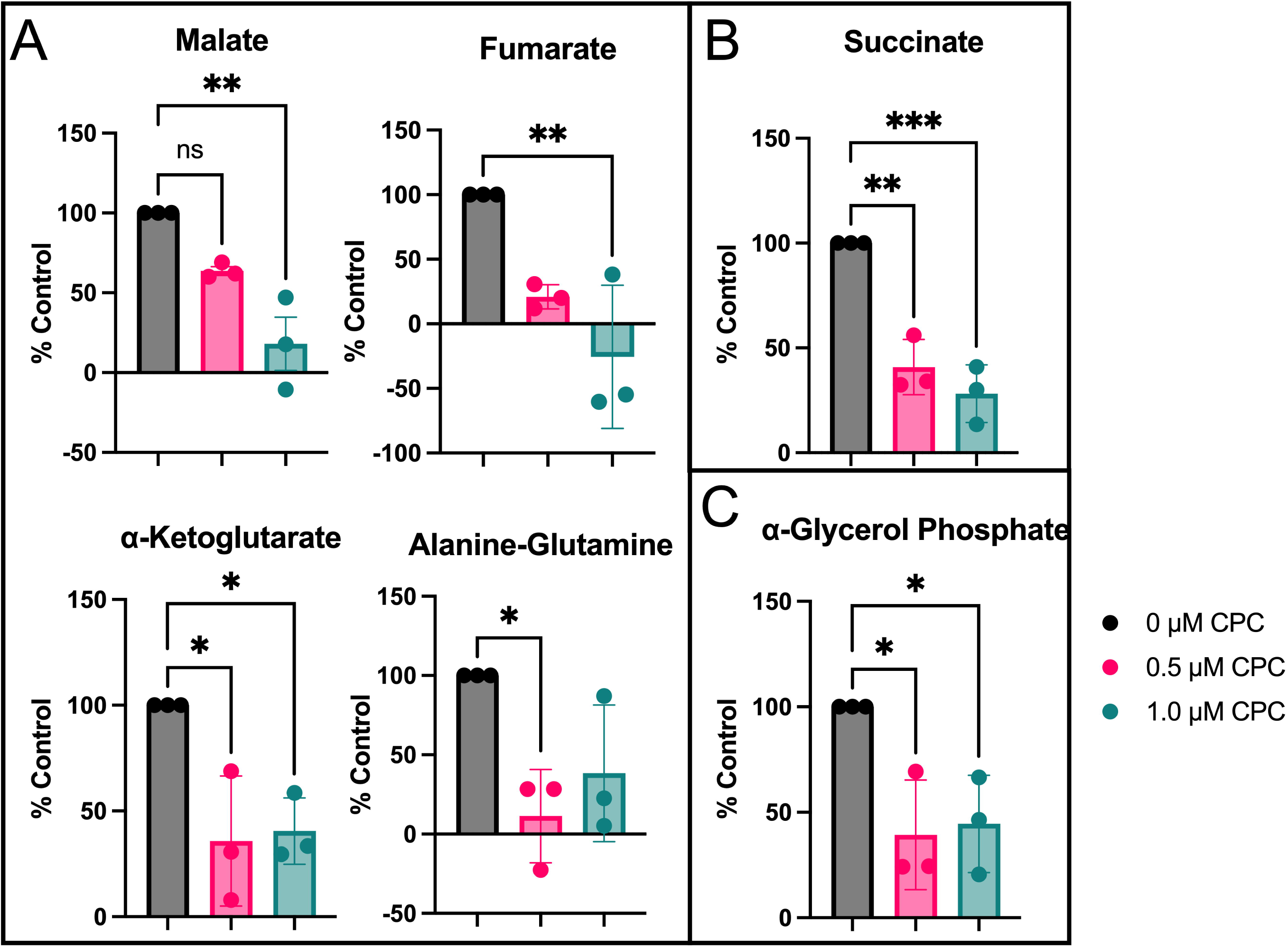
Effect of CPC on electron flow through the mitochondrial electron transport chain in primary human keratinocytes. Electron flow was measured in saponin-permeabilized primary human keratinocytes (via A_590_, plate reader) from several metabolic substrates through cytochrome c during an up to 3 h exposure (details in Table S2) to control buffer, versus 0.5 μM or 1.0 μM CPC (Figure 1). The y-axis represents the rate of electron flow for each individual substrate, normalized to the no-substrate control of each given day. A t-test was used (Fig. S1B) to determine which of the 31 substrates produced significant electron flow compared to the no-substrate control. Substrates (6) passing the t-test were grouped, for display, by entry point into the chain: complex I-feeding substrates (**A**), complex II-feeding substrate succinate **(B**), and CoQ-feeding substrate α-glycerol phosphate (**C**). Data shown are mean ± SEM. Data gathered from 3 days of experiments. Significant results determined by one-way ANOVA and Dunnett’s post-test (95% CI). ***p < 0.001, **p < 0.01, *p < 0.05.

**Figure 3.**
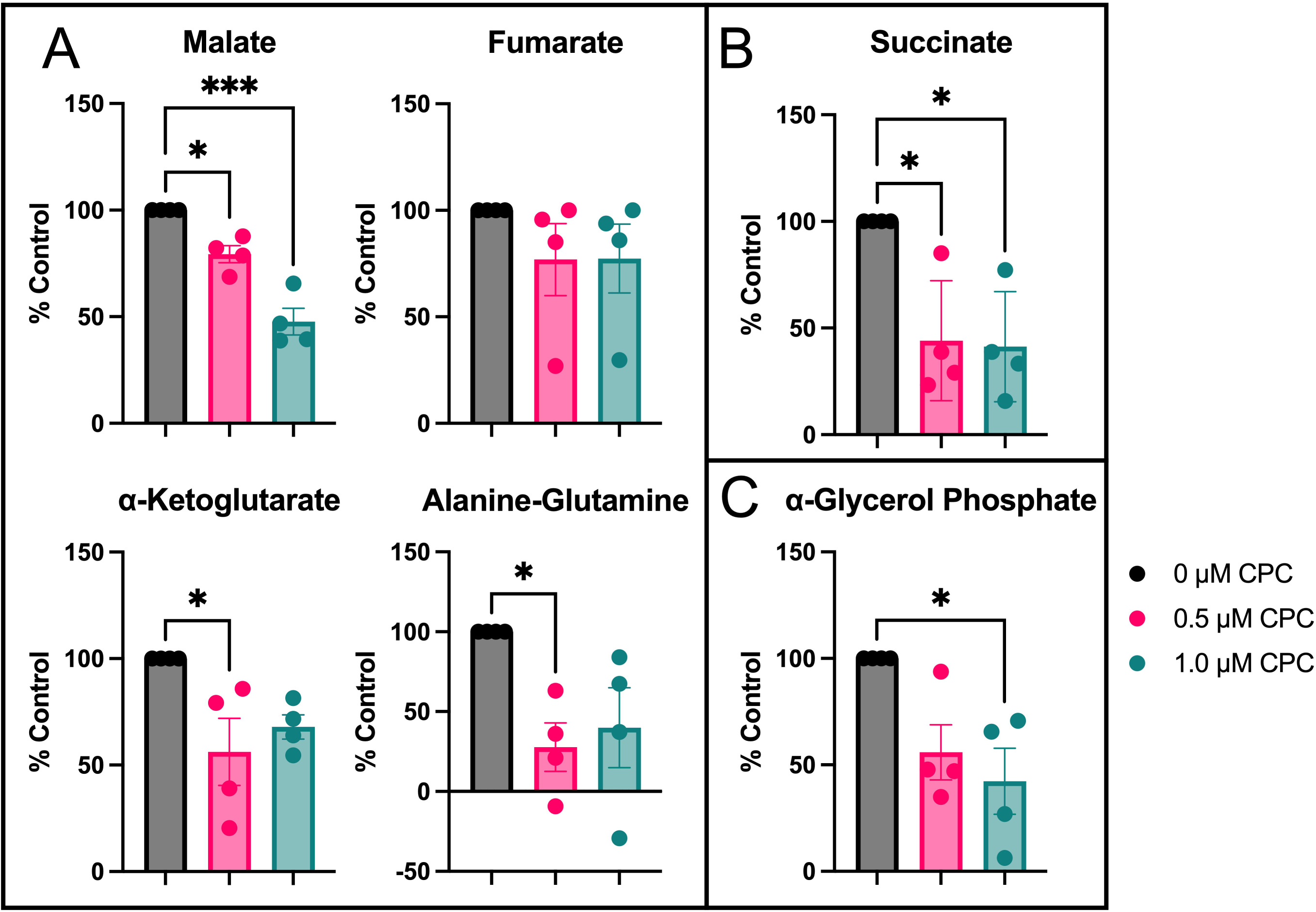
Effect of CPC on electron flow through the mitochondrial electron transport chain in RBL-2H3 mast cells. Electron flow was measured in saponin-permeabilized RBL-2H3 mast cells (via A_590_, plate reader) from several metabolic substrates through cytochrome c during an up to 2 h exposure (details in Table S2) to control buffer, 0.5 μM or 1.0 μM CPC (Fig 1). The y-axis represents the rate of electron flow for each individual substrate, normalized to the no-substrate control of each given day. A t-test was used (Fig. S2B) to determine which of the 31 substrates produced significant electron flow compared to the no-substrate control. Substrates (6) passing the t-test were grouped, for display, by entry point into the chain: complex I-feeding substrates (**A**), complex II-feeding substrate succinate (**B**), and CoQ-feeding substrate α-glycerol phosphate (**C**). Data shown are mean ± SEM. Data gathered from 4 days of experiments. Significant results determined by one-way ANOVA and Dunnet’s post-test (95% CI). ***p < 0.001, *p< 0.05.

**Figure 4.**
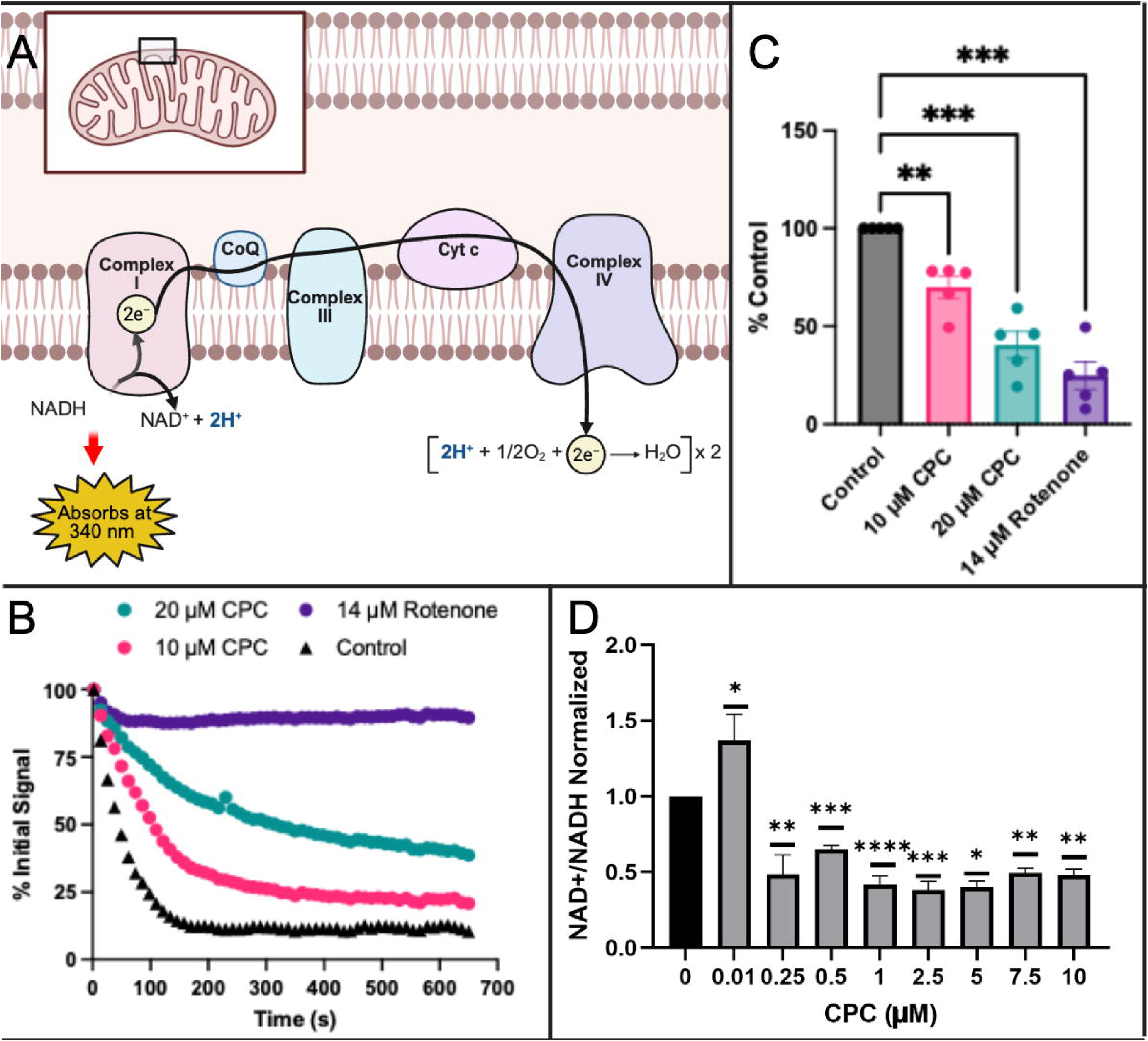
Effect of CPC on electron flow from NADH through the mitochondrial electron transport chain in isolated bovine mitochondria and in intact RBL-2H3 mast cells. Mitocheck Complex I colorimetric assay in isolated mitochondria and Promega NADH-Glo kit in intact mast cells were used to assess electron flow from NADH through the full chain. For each experiment, mitochondria or cells were pre-exposed for 1 h to the noted CPC or rotenone treatments. (**A**) Schematic showing Mitocheck Complex I assay, in which mitochondria were fed NADH to start the reaction, and NADH was also the probe (read via A_340_, plate reader). Made in Biorender. (Cyt C, cytochrome c; CoQ, coenzyme q). (**B**) Mitocheck Complex I data of NADH oxidation, graphed as percentage of initial signal over time, averaged over 5 days of experiments (duplicates per experiment), following subtraction of background signal (from mitochondria in the buffer but with no added NADH). (**C**) Slopes from the first 98 sec of data in Fig. 4B, normalized to the no-treatment control of each given day, are plotted to visualize and compare treatment effects on NADH oxidation. Data are displayed as mean ± SEM. (**D**) A luminescence-based NADH-Glo assay was used to measure the NAD+/NADH ratio in intact RBL-2H3 cells. Data gathered from 4 experiments, 3 replicates per experiment, are shown as mean ± SEM. For **C** and **D**, significance analysis was performed via one-way ANOVA with Dunnett’s post-hoc test, ***p < 0.001, **p < 0.01, *p < 0.05.

*Complex I Data Analysis:* To analyze the experiments, it was necessary to first determine the linear range. For these experiments, the “linear range” was defined by the longest interval at which the R^2^ value (obtained using the RSQ Microsoft Excel function) was > 0.9 for the entire data set as a whole (Fig. S3C). Consequently, the timeframe used for analysis was 98 s (Fig. S3C). These timepoints also maximized the amount of signal range: starting absorbance values were robust, with A_340_ ranging from 0.134 to 0.241 over all experimental days. Additionally, the dynamic range of the experiments used in Fig. 4C was from 0.113 to 0.209, a robust change with capacity to detect either increases or decreases in Complex I activity. After determination of the linear range, the slope at the chosen timepoint (98 s) was found using Excel for each condition, then normalized to the control on that day. Replicates were put into GraphPad Prism and analyzed with a one-way ANOVA and Dunnett’s multiple comparisons test to the 0 µM CPC data.

## NADH-Glo Assay

This assay utilized the NAD/NADH-Glo™ kit from Promega, and the protocol was per manufacturer’s instructions, with added adaptations from(Gui et al., 2016). RBL-2H3 mast cells were plated in glucose-free galactose-DMEM media(Weatherly et al., 2016) at 50,000 cells per well and left to adhere for 3 hours. CPC concentrations were made up in glucose-free galactose-DMEM media with BSA (1 mg/mL). Cells were then exposed to CPC solutions for 60 min. Cells were lysed with a 1% (w/v) solution of dodecyltrimethylammonium bromide in 0.2 N NaOH, diluted to 0.5% (w/v) with PBS. Lysates were mixed and split into two samples: a 25 µL sample (for NADH analysis) and 20 µL sample (for NAD+ analysis). Within each NAD+ selection tube, there was an additional 20 µL of lysis buffer and 20 µL of 0.4 N HCl. Within each NADH selection tube, there was no additional solution. NADH tubes were heated to 75°C for 30 min, and NAD+ tubes were heated to 60°C for 15 min. Tubes were brought to room temperature for 8 min, then quenched with 25 µL 0.25 M Tris in 0.2 N HCl for NADH tubes, and in 20 µL 0.5 M Tris for NAD+ tubes. Mixed samples of 50 µL from each tube were then moved to a white-bottom 96-well plate and each well additionally contained 50 µL of NAD/NADH-Glo™ Detection Reagent (Promega) per manufacturer’s instructions. The plate was incubated at room temperature for 30 min, then read for luminescence (Synergy H1 plate reader, Agilent Biotek). For data analysis, wells were background-subtracted (using background samples containing all assay components other than cells or CPC), and averaged. Replicates were put into GraphPad Prism and analyzed with a one-way ANOVA and Dunnett’s multiple comparisons test to the 0 µM CPC data.

## Mitocheck Complex II

### Preparation of Antimycin A

Antimycin A (Sigma Aldrich; CAS No. 1397-94-0) i is virtually insoluble in water (Pubchem). Therefore, antimycin A was first dissolved into 100% DMSO at 10 mM and diluted in Complex II buffer. Based on the ultra-violet spectra obtained from supplier Cayman Chemical, Antimycin A does not interfere with any of the wavelengths used in the corresponding MitoCheck assays (Table S3). Vehicle controls were performed to ensure that the DMSO vehicle did not interfere with the Complex III assay as well since it required Antimycin A (Fig. S5B).

### Preparation of Thenoyltrifluoroacetone

Thenoyltrifluoroacetone (TTFA; Sigma-Aldrich, CAS No. 326-91-0) is soluble in water up to only 0.34 mM (Pubchem). Therefore, TTFA was dissolved in 100% DMSO, making a concentration of 120 mM which was diluted to a final 1X concentration of 2.5 mM TTFA in the well with all assay constituents. Based on the ultra-violet spectra obtained from supplier Cayman Chemical, TTFA does not interfere with the Complex II readout measurement (which is 600 nm) (Table S3). Vehicle controls were performed to ensure that the DMSO vehicle did not interfere with the assay (Fig. S4B).

### Mitocheck Complex II Data Collection

MitoCheck Complex II Activity assay kit (Cayman Chemical) measures electron flow from succinate through Complex II, CoQ, and to the dichlorophenolindophenol (DCPIP) dye (Fig. 5A). All materials used except for the treatment conditions were provided with the assay kit. Isolated bovine heart mitochondria were dissolved into Complex II buffer. Inhibitors rotenone (2 uM) and Antimycin A (14 uM) were added to isolate electron flow to the Complex II route of the ETC. The mitochondria (in assay solution with inhibitors) were then pre-exposed to each treatment group (CPC, positive control inhibitor TTFA, or vehicle control) for 60 min at 37 °C/5% CO_2_. Following the aforementioned 60 min pre-treatment, succinate, CoQ, and DCPIP dye were added per manufacturer’s instructions. Complex II activity was then measured in MitoCheck Complex II buffer with DMSO (0.14%) as required by the TTFA dissolution, in order to standardize the vehicle across all treatment types. Complex II enzyme activity, correlating to the rate of DCPIP dye reduction, was measured via kinetic A_600_ absorbance reading in the plate reader (Synergy H1 plate reader, Agilent Biotek) for 20 min at 12 s intervals and at 25 °C. Example background-subtracted results are shown in Fig. S4A.

**Figure 5.**
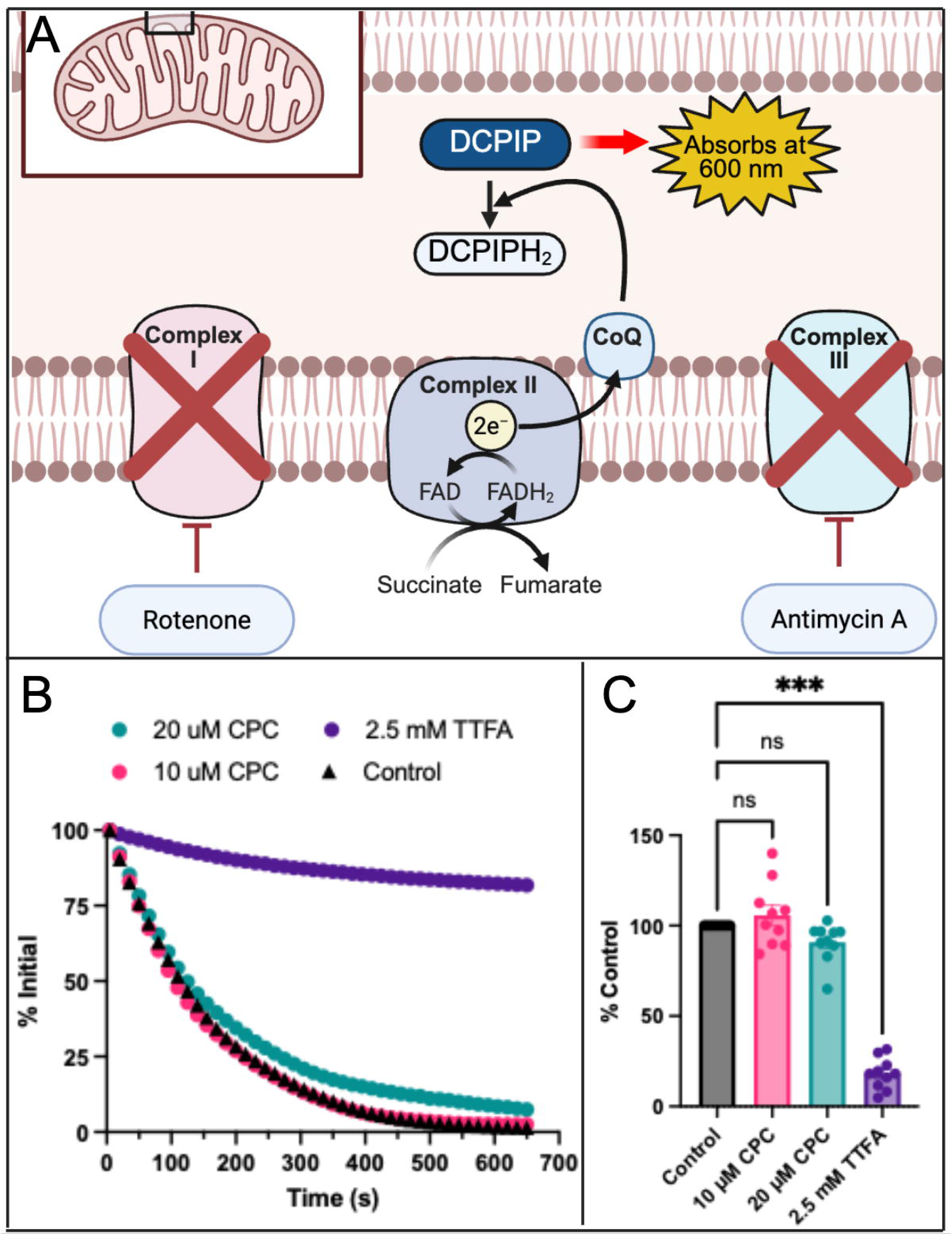
Effect of CPC on electron flow from succinate through Complex II and CoQ in isolated bovine mitochondria. (**A**) Mitocheck Complex II colorimetric assay in isolated mitochondria was used to assess electron flow from succinate through CoQ to a DCPIP redox dye. For each experiment, mitochondria were pre-exposed for 1 h to the noted CPC or TTFA treatments. For each sample, mitochondria were simultaneously exposed to rotenone to inhibit Complex I and to antimycin A to shut down Complex III, so as to isolate Complex II for investigation. (**A**) Schematic showing Mitocheck Complex II assay, in which mitochondria were fed succinate to start the reaction, supplied with excess CoQ in order to focus analysis on Complex II, and DCPIP was the probe (read via A_600_, plate reader); the 2 inhibitors are also indicated. Made in Biorender. (CoQ, coenzyme q; DCPIP, 2,6-dichlorophenolindophenol; TTFA, thenoyltrifluoroacetone). (**B**) Mitocheck Complex II data of DCPIP oxidation, graphed as percentage of initial signal over time, averaged over 9 days of experiments (duplicates per experiment), following subtraction of background signal (from mitochondria in the buffer but with no added DCPIP dye). (**C**) Slopes from the first 86 sec of data in Fig. 5B, normalized to the no-treatment control of each given day, are plotted to visualize and compare treatment effects on DCPIP oxidation. Data are displayed as mean ± SEM. Significance analysis was performed via one-way ANOVA with Dunnett’s post-hoc test, ***p < 0.001, ns= not significant.

### Complex II Data Analysis

To analyze the experiments, it was necessary to first determine the linear range. For these experiments, the “linear range” was defined by the longest interval at which the R^2^ value (obtained using the RSQ Microsoft Excel function) was > 0.95 for the entire data set as a whole (Fig. S4C). Consequently, the timeframe used for analysis was 86 s (Fig. S4C). These timepoints also maximized the amount of signal range: starting absorbance values were robust, with A_600_ ranging from 0.302 to 1.230 over all experimental days. Additionally, the dynamic range of the experiments used in Fig. 5C was from 0.302 to 1.2963, a robust change. After determining the linear range, the slope at the chosen timepoint (86 s) was found using Microsoft Excel for each condition, then normalized to the control on that day. Replicates were put into GraphPad Prism and analyzed with a one-way ANOVA and Dunnett’s multiple comparisons test to the 0 µM CPC data.

## Mitocheck Complex III

*Preparation of potassium cyanide (KCN):* In assays of Complex III and IV, KCN (CAS No 151-50-8; Sigma-Aldrich) was utilized. KCN was prepared in basic buffer with NaOH to prevent protonation of the KCN. It is well-established that cyanide is largely inhibitory against Complex IV when in its ionic form (cyanide as opposed to the protonated hydrogen cyanide form)(Lachowicz et al., 2024; Leavesley et al., 2008; Nelson & Cox, 2013). NaOH (CAS No. 310-73-2; VWR) was diluted from 10 M to 0.05 M with CCW. KCN, which is highly soluble in water at 11 M (Pubchem), was dissolved into 0.05 M NaOH to make 10 mM KCN stock solution. Complex IV experiments with NaOH were conducted in Mitocheck buffer at pH 7.8 (following the addition of the KCN in NaOH), higher than physiological but required for cyanide activity). This pH was chosen in order to be close enough to physiological for sufficient mitochondrial activity: In Fig. S6B, we show that this higher-pH buffer does not change the activity of Complex IV significantly. At this pH, the Henderson-Hesselbach equation (using cyanide’s pK_a_ 9.21) indicates that only ∼4% of the cyanide molecules are in their anionic form, the form effective against Complex IV. Thus, we were required to use 25X more KCN than is stated in Fig. 7. For example, Fig. 7 states that 10 uM cyanide was used, but this is the calculated effective CN^-^ concentration while the total made KCN concentration was actually 250 uM; this situation is the same in Complex III experiments (Fig. 6).

**Figure 6.**
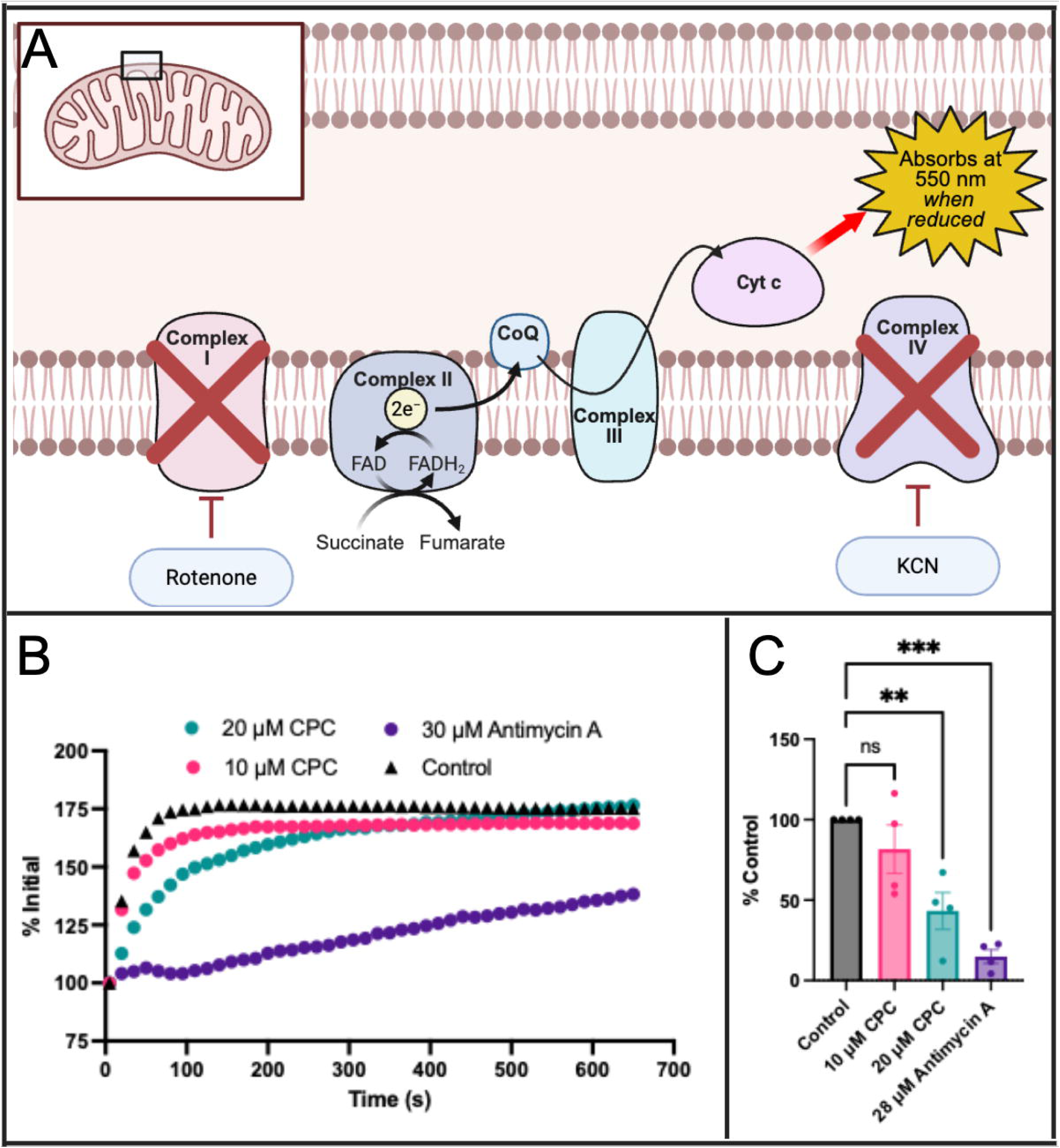
Effect of CPC on electron flow from succinate through cytochrome c in isolated bovine mitochondria. Mitocheck Complex III colorimetric assay in isolated mitochondria was used to examine electron flow from succinate through complex III to cytochrome c. For each experiment, mitochondria were pre-exposed to CPC or Antimycin A for 1 h. For each sample, mitochondria were simultaneously exposed to rotenone to inhibit Complex I and to KCN to shut down Complex IV, so as to isolate Complex III and Cyt c for investigation. (**A**) Schematic showing Mitocheck Complex III assay, in which mitochondria were fed succinate to start the reaction, and Cyt c was the probe (read via A_550_, plate reader); the 2 inhibitors are also indicated. Made in Biorender. (CoQ, coenzyme q; Cyt c, cytochrome c). (**B**) Mitocheck Complex III data of Cyt c reduction, graphed as percentage of initial signal over time, averaged over 4 days of experiments (duplicates per experiment), following subtraction of background signal (from mitochondria in the buffer but with no added Cyt c). (**C**) Slopes from the first 50 sec of data in Fig. 6B, normalized to the no-treatment control of each given day, are plotted to visualize and compare treatment effects on Cyt c reduction. Data are displayed as mean ± SEM. Significance analysis was performed via one-way ANOVA with Dunnett’s post-hoc test, ***p < 0.001, **p<0.01.

**Figure 7.**
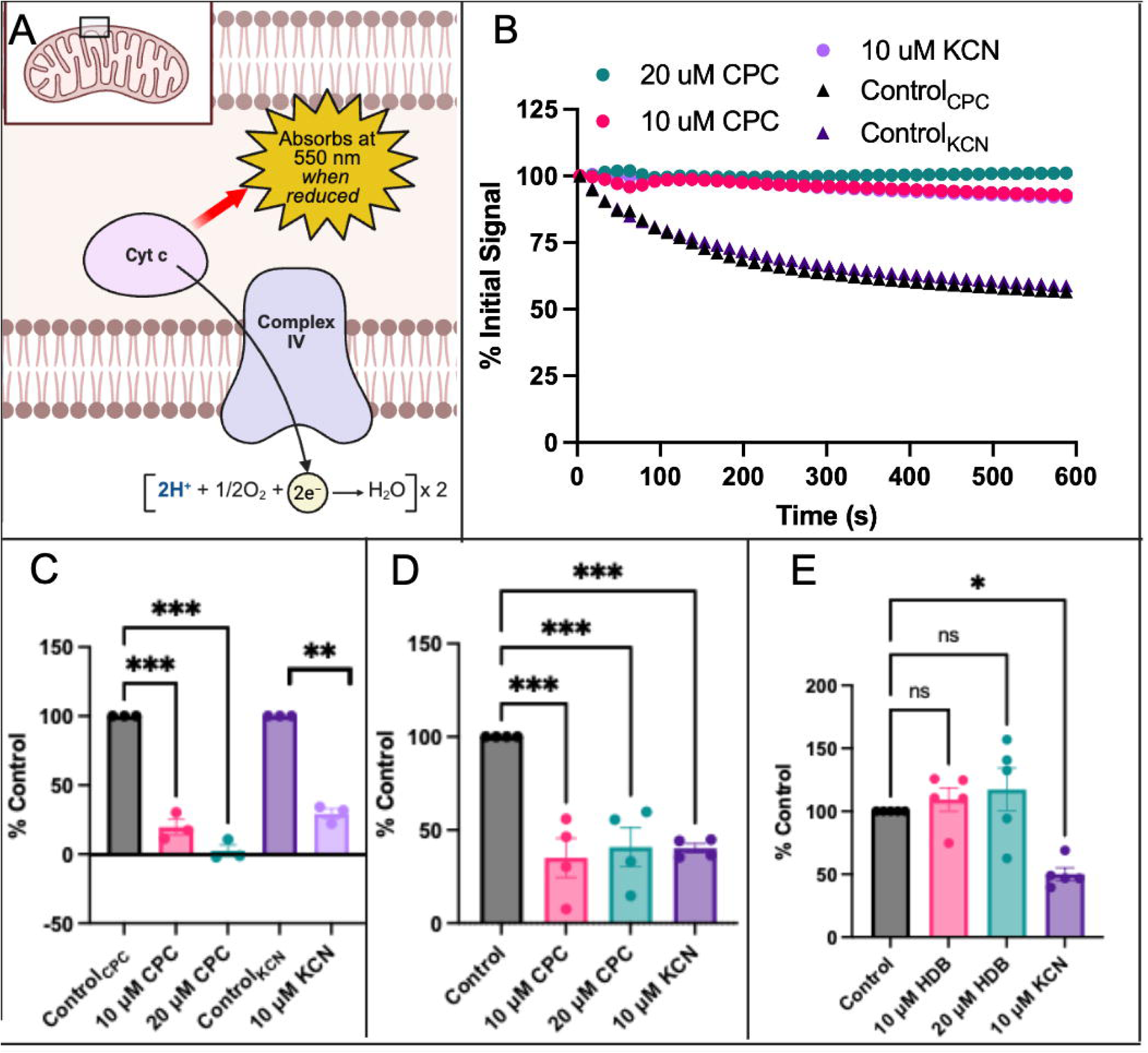
Effect of CPC on electron flow from cytochrome c to Complex IV in isolated bovine mitochondria. Mitocheck Complex IV assay in isolated mitochondria was used to analyze electron flow from Cyt c to Complex IV. For each experiment, mitochondria were pre-exposed to CPC, KCN, or HDB for 1 h. **A**) Schematic showing Mitocheck Complex IV assay, in which mitochondria were fed Cyt c to start the reaction, and Cyt c was also the probe (read via A_550_ plate reader). Made in Biorender. (Cyt c, cytochrome c). (**B**) Mitocheck Complex IV data of Cyt c oxidation, graphed as percentage of initial signal over time, averaged over 3-5 days of experiments (duplicates per experiment), following subtraction of background signal (from mitochondria in the buffer but with no added Cyt c). (**C**) Slopes from 3-10 min of data in Fig. 7B, normalized to each separate no-treatment control, are plotted to visualize and compare treatment effects on Cyt c reduction. (**D**) Slopes from 3-10 min of data from CPC experiments in common NaOH buffer, normalized to the no-treatment control of each given day. (**E**) Slopes from 3-10 min of data from HDB experiments in common NaOH buffer, normalized to the no-treatment control of each given day. Data in each graph are displayed as mean ± SEM. Significance analysis was performed via one-way ANOVA with Dunnett’s post-hoc test, ***p < 0.001, **p < 0.01, *p<0.05, ns= not significant.

*Mitocheck Complex III Data Collection:* MitoCheck Complex III Activity assay kit (Cayman Chemical) measures electron flow from succinate through Complex II, then CoQ, Complex III, and finally to cyt C (Fig. 6A). All materials used except for the treatment conditions were provided with the assay kit. Isolated bovine heart mitochondria were dissolved into Complex III buffer with inhibitors rotenone (2 uM) and KCN (see above) which were added to isolate electron flow through Complex III. The mitochondria were then pre-exposed to each treatment group (CPC, positive control inhibitor Antimycin A, or vehicle control) for 60 min at 37 °C/5% CO_2_.

Complex III activity was then measured in MitoCheck Complex III buffer with DMSO (0.28%) as required by the Antimycin A dissolution, in order to standardize the vehicle across all treatment types. Following the aforementioned 60 min pre-treatment, succinate, CoQ, and oxidized cyt C were added per manufacturer’s instructions.

Complex III enzyme activity, correlating to the rate of cyt C reduction, was measured via kinetic A_550_ absorbance reading in the plate reader (Synergy H1 plate reader, Agilent Biotek) for 20 min at 12 s intervals and at 25 °C. Example background-subtracted results are shown in Fig. S5A.

*Complex III Data Analysis:* To analyze the experiments, it was necessary to first determine the linear range. For these experiments, the “linear range” was defined by the longest interval at which the R^2^ value (obtained using the RSQ Microsoft Excel function) was > 0.80 for the entire data set as a whole (Fig. S5C). Consequently, the timeframe used for analysis was 50 s (Fig. S5C). These timepoints also maximized the amount of signal range: starting absorbance values were robust, with A_550_ ranging from 0.087 to 0.214 over all experimental days. Additionally, the dynamic range of the experiments used in Fig. 6C was 0.084-0.195 absorbance units, a robust change. After determination of the linear range, the slope at the chosen timepoint (50 s) was found using Microsoft Excel for each condition, then normalized to the control on that day. Replicates were put into GraphPad Prism and analyzed with a one-way ANOVA and Dunnett’s multiple comparisons test to the 0 µM CPC data.

## Mitocheck Complex IV

*Preparation of hexadecylbenzene*: In addition to CPC testing, hexadecylbenzene (HDB) was utilized in this assay. HDB, which is structurally similar to CPC but lacks CPC’s positive head group, was prepared as in(Obeng et al., 2024), with the exception that it was dissolved into Complex IV Mitocheck buffer.

*Mitocheck Complex IV Data Collection:* MitoCheck Complex IV Activity assay kit (Cayman Chemical) measures electron flow from cyt C to Complex IV (Fig. 7A). All materials used except for the treatment conditions were provided with the assay kit. Isolated bovine heart mitochondria were dissolved into Complex IV buffer then were pre-exposed to each treatment group (CPC, positive control inhibitor KCN, HDB, or vehicle control) for 60 min at 37 °C/5% CO_2_. For one set of experiments (Fig. 7C), CPC and KCN were tested in separate buffers: CPC in “Control_CPC_” MitoCheck Complex IV buffer was tested concurrently to KCN, which was in the same buffer plus added 0.05 M NaOH (“Control_KCN_” buffer) (these are labeled in Fig. 7C). For the other set of experiments (Fig. 7D), all samples tested (CPC or KCN) were dissolved in the same 0.05 M NaOH-containing Complex IV buffer, which contained an identical amount of NaOH across all treatment groups, done to allow for direct comparison of CPC with KCN (this is simply called “Control” in Fig. 7D). Note that this “Control” (or “Control_KCN_”) Buffer was calculated to be pH ∼7.8 due to the added NaOH. For the HDB analysis, this “Control” Buffer was used, for which HDB and KCN were both dissolved in the same 0.05 M NaOH-containing Complex IV buffer.

Complex IV activity was then measured. Following the aforementioned 60 min pre-treatment, reduced cyt C was added per manufacturer’s instructions. Complex IV enzyme activity, correlating to the rate of cyt C oxidation, was measured via kinetic A_550_ absorbance reading in the plate reader (Synergy H1 plate reader, Agilent Biotek) for 20 min at 15 s intervals and at 25 °C. Example background-subtracted results are shown in Fig. S6A.

*Complex IV Data Analysis:* To analyze the experiments, it was necessary to first determine the linear range. For these experiments, the “linear range” was defined by the longest interval at which the R^2^ value (obtained using the RSQ Microsoft Excel function) was > 0.9 for the entire data set as a whole (Fig. S6C). Consequently, the timeframe used for analysis was 3-10 min (Fig. S6C). These timepoints also maximized the amount of signal range: starting absorbance values were robust, with A_550_ ranging from 0.2621 to 0.425 over all experimental days. Additionally, the dynamic range of the experiments used in Fig. 7C was from 0.113-0.24, and the range of experiments in Fig. 7D and 7E (all in the NaOH-containing buffer) were 0.031-0.098. After determination of the linear range, the slope at the chosen timepoint (3-10 min) was found using Microsoft Excel for each condition, then normalized to each corresponding control on that day. Replicates were put into GraphPad Prism and analyzed with a one-way ANOVA and Dunnett’s multiple comparisons test to controls.

## Assay of CPC Effect on the Redox State of Coenzyme Q10 in Cells

RBL-2H3 cells were plated in a 6-well clear-bottom tissue culture-treated (2 million cells/well) in phenol red-free RBL media and allowed to attach for ∼12-18 hours.

The next day, glucose-BSA-Tyrodes (glucose-BT) was prepared by dissolving bovine serum albumin (BSA, 1 mg/mL) in Tyrodes solution at pH 7.4(Hutchinson et al., 2011). CPC was prepared in glucose-BT. The cells were then rinsed with glucose-BT and treated for 1 hour in glucose-BT (control) or 5 or 10 uM CPC doses (in glucose-BT) at 37 °C/5% CO_2_. The cells were then washed twice in glucose-BT, wash was discarded, and 0.3 mL PBS was added per well. Cells were scraped and pipetted into 1.5 mL tubes, then lysed by sonication: 20 % power, sonicate 3 s on, 10 s off, then repeated 30 times (Branson SFX 250 Sonifier; Emerson Electric). To standardize the cellular contents per well of each experimental group, we used the Bradford protein assay (VWR).

A colorimetric Coenzyme Q10 (CoQ) assay (Arigo) was then used to determine levels of reduced CoQ in treated versus control cells. Assay buffer (0.3 mL) was added to each sonicated sample, which were then centrifuged at 10000xg for 10 min at 40 °C. A colorimetric dye was employed to detect CoQ via A_620_ (plate reader, Synergy H1 plate reader, Agilent Biotek). A CoQ standard curve was made using a standard of pure CoQ. From the standard curve, CoQ levels were calculated for CPC versus control levels. Statistical analysis was done in Graphpad Prism using one-way ANOVA and Dunnet’s post-hoc test.

To ensure these data were reproducible in cells fed glucose-free galactose DMEM, these methods were additionally performed in glucose-free galactose-DMEM media.

## Total Cardiolipin Level Determination

CPC effects on cardiolipin (CL) levels within cells were probed with a cardiolipin assay kit (Abcam). RBL-2H3 cells were cultured as previously described and seeded into a 6-well plate at 2,000,000 cells/well in 3 mL phenol red-free RBL media for 16-18 h.

The next day, cells were washed with glucose-BT twice, then treated with either 1 mL/well of 0, 5, or 10 μM CPC (prepared in glucose-BT) for 1 h at 37 °C/5% CO_2_. After treatment, cells were washed with PBS (2 mL/well) twice and scraped (using cell scrapers) into Abcam assay buffer (500 uL/well) and each placed into 1.5 mL microcentrifuge tubes. Cell samples were mechanically lysed via probe sonication using a Branson SFX 250 Sonifier (Emerson Electric) (50 % power, sonicate 1 s followed by 3 s off, 3 repeats of this cycle). Afterwards, lysates were centrifuged at 4 °C for 10 minutes at 10,000 xg. The Bradford protein assay (VWR) was conducted on the supernatant to ensure equal amounts of cellular contents were assessed for each sample. Assays were performed according to manufacturer’s instructions and measured in the plate reader (Synergy H1 plate reader, Agilent Biotek). Known standards of cardiolipin were then prepared using Abcam’s CL kit to construct a standard curve, to calculate the amount of CL present in each cell sample. For each experiment, two background controls were conducted and then summed together, to act as the background: 1.) a probe background (containing assay buffer and probe) and 2.) a control cell lysate (assay buffer and untreated but lysed cells). Each background subtracted treatment condition was normalized to the untreated-cell (0 uM CPC) CL value for each day. A one-way ANOVA with Dunnett’s post-hoc was conducted using GraphPad Prism.

To ensure these data were reproducible in cells fed glucose-free galactose DMEM, these cardiolipin level methods were also performed in glucose-free galactose-DMEM media (**Fig. S7**).

## Cardiolipin Dye Experiment

CPC effects on purified cardiolipin’s ability to bind its probe were investigated with the same CL assay kit (Abcam). CPC was prepared in CCW. Purified CL was reconstituted to 5 mM in 100% ethanol. CL solutions were prepared at 0, 1, and 10 uM concentrations in kit Assay buffer with various CPC doses, such that the 0.2% ethanol vehicle was present. These solutions were incubated for 15 min, shaking at 150 rpm (Gyrotary Shaker Model G2, New Brunswick Scientific Company) at room temperature. Fluorescence of Abcam CL probe was then detected via plate reader (Synergy H1, Agilent Biotek) per manufacturer’s instructions. Probe fluorescence was normalized to the maximum signal intensity (10 uM cardiolipin, no CPC) and graphed against cardiolipin concentration for each CPC condition (0, 5, and 50 uM CPC). Three experiments were carried out. A one-way ANOVA with Dunnett’s post-hoc was conducted using GraphPad Prism.

## Cardiolipin Bead Assay

*Preparation of 1-methylpyridinium chloride:* 1-methylpyridinium chloride (MPC; CAS No. 7680-73-1, 98% purity, Tokyo Chemical Industries) was prepared in the same manner as CPC, dissolved in CCW and sonicated. Both CPC and MPC were checked for concentration in UV-Vis as described(Raut, Weller, et al., 2022).

*Cardiolipin bead assay development*: We developed a novel assay using cardiolipin-coated beads (Echelon) to assess the effect of CPC on CL binding to its known binding partner cyt C. This assay requires the use of phosphate buffer (pH 7, CAS No 7558-79-4; Thermo Fisher), including 0.1% TWEEN-20 (Sigma) to reduce nonspecific interactions. Published work indicates that Tween detergents may drastically reduce the critical micelle concentration of CPC, drawing it into micelles(Ghosh, 2001) in such a way that might reduce its ability to directly bind lipids like CL. To avoid this complication, we instead employed MPC, which is the positively-charged head group from CPC, capped by a methyl group in place of the lipid tail.

Thus, we are testing the role of CPC’s charged head group without the uncharged lipid tail with which TWEEN-20 would interfere. Additionally, Echelon called for use of a high NaCl concentration, like that in blood/extracellular levels and actually far higher than [NaCl] in the cytosol/intermembrane space. However, high salt is well-known to disrupt protein-lipid interactions, so instead we used a mitochondrially-relevant NaCl concentration of 10 mM(Baggaley & Riedel, 1966), to allow for CL-cyt C binding.

*Detection of MPC effect on CL bead binding to purified cytochrome:* Echelon’s CL-coated bead slurry was well-mixed and transferred into experimental 0.6 mL tubes (50 uL/sample). Storage buffer was removed, and 50 uL/tube phosphate buffer (with 10 mM NaCL, 0.1% Tween-20) was added. Beads were then pre-exposed to MPC (0 control, 10, or 20 uM) for 1 h at room temperature with agitation (Gyrotary Shaker Model G2, New Brunswick Scientific Company), with manual inversion of the tubes every 15 min. The supernatant was then discarded. Next, 50 uL of the same MPC solutions were added to each respective sample, with the addition of 2 ug reduced cyt C (equine heart; Cayman Chemical), and the mixtures were incubated at room temperature for 3 h with agitation (Gyrotary Shaker Model G2, New Brunswick Scientific Company). Every 15-30 min during the 3 h, tubes were manually inverted. (Note: while both reduced and oxidized cyt C forms bind to CL(Milorey et al., 2016), we utilized the reduced form to best simulate the electron transfer to Complex IV.)

The supernatant from each experimental tube (0 control, 10, or 20 uM MPC) was removed and plated into a clear half-area microplate (VWR), then read for A_410_ (cyt C’s Soret band, (Margoliash & Frohwirt, 1959) via plate reader (Synergy H1, Agilent Biotek). Supernatant absorbance represents cyt C that was not bound to CL at the time assayed, based upon published methods(Caesar et al., 2009). An increase in A_410_ signal denotes MPC disruption of cyt C’s ability to bind CL. Data were normalized to control on each day, and statistically tested with one-tailed, nonparametric t-test.

## Statistical Analysis

All analyses were performed using Prism software (Graphpad, San Diego, CA). Means ± SEM were determined by averaging biological replicates from at least three independent days of experiments. Specific statistical tests employed for each assay type are described in the respective sections.

## Results

*Characterization of CPC-Induced Mitochondrial Metabolic Phenotype via Biolog MitoPlate S-1.* In the presence of saponin permeabilization of the plasma membrane, the Biolog MitoPlate S-1 96-well format can be used to screen mitochondrial metabolism from multiple separate purified individual substrates. Fig. 1 details the path of electron flow from these various substrates (TCA cycle intermediates, ⍰-glycerol phosphate), via NADH or FADH_2_, through Complexes I-III of the ETC, then to the Biolog dye (MC redox dye) after cytochrome c (cyt C). Notably, Complex IV is not assayed by this method: thus, the source of any observed effects on electron flow must be upstream of Complex IV (between the individual substrates and cyt C). These experiments are performed in minimal-nutrient MAS buffer; thus, the vast majority of electrons are provided by each individual test substrate.

Low CPC doses were tested on electron flow in the mitochondria of primary human keratinocyte (Fig. 2) and RBL-2H3 cells (Fig. 3). To focus on mitochondrial effects, low level saponin is used to permeabilize the plasma membrane without harming the mitochondrial membranes. Thus, some of the plate’s substrates cannot be metabolized (e.g. sugars that require intact cytosolic glycolysis). Successful electron flow from each substrate through cyt C is represented by MC redox dye reduction, resulting in an increase in A_590_ signal over time (electron flow rate). (Analyses confirmed that dye signal is unaffected by CPC exposure, Fig. S1). Of the substrates assayed under saponin permeabilization conditions, six induced statistically-significant electron flow (Table S1): malate, ⍰-ketoglutarate, fumarate, alanine-glutamine dipeptide (Ala-Gln), succinate, and ⍰-glycerol phosphate. Each of these is a mitochondrially-processed electron source.

Complex I collects electrons from the carrier NADH, which is produced by TCA enzymes that process the intermediates malate and ⍰-ketoglutarate. Fumarate also leads to NADH/Complex I because fumarase creates malate. Furthermore, the test substrate Ala-Gln is mitochondrially converted to ⍰-ketoglutarate/pyruvate; therefore Ala-Gln also feeds NADH to Complex I, although indirectly. Of these four Complex I-feeding substrates (malate, ⍰-ketoglutarate, fumarate, and Ala-Gln), low-micromolar CPC significantly inhibits their electron flow in both primary human keratinocytes (Fig. 2A) and RBL-2H3 mast cells (Fig. 3A, with the exception of fumarate in the RBLs). CPC modesty inhibits electron flow rate from fumarate in RBLs, but the effect only reached the level of significance in primary human keratinocytes. Electron flow rate from malate and ⍰-ketoglutarate is decreased by ≥ 50% (in both cell types), due to 0.5 or 1 uM CPC. Electron flow from Ala-Gln is decreased by ∼75% (in both cell types) by 0.5 uM CPC.

Complex II (succinate dehydrogenase), both a member of the TCA cycle and the mitochondrial electron transport chain, collects electrons from the intermediate succinate, via carrier FADH_2_. Electron flow rate from succinate to the MC redox dye (Fig. 2B, 3B) is also significantly inhibited by 0.5 uM CPC in both cell types, by ≥ 50%.

Electrons enter into CoQ of the ETC from the intermediate ⍰-glycerol phosphate, via ⍰-glycerol phosphate dehydrogenase using carrier FADH_2_, circumventing both Complexes I and II. Electron flow rate from ⍰-glycerol phosphate (Fig. 2C, 3C) is also significantly inhibited by 0.5 uM CPC in both cell types, by ≥ 50%.

Taken together, the Biolog MitoPlate S-1 results show CPC inhibits electron flow at some point after each of these aforementioned substrates, yet before the dye intercedes, just after cyt C.

*CPC Does Not Directly Affect TCA Enzyme Malate Dehydrogenase*. Because malate consistently displayed the strongest electron flow signal in the Biolog assay as well as robust CPC effects, we investigated whether CPC directly inhibits mitochondrial malate dehydrogenase 2 (MDH2). Supplement Fig. S2 describes this work and shows that CPC does not directly inhibit this TCA enzyme, pointing toward the electron transport chain as CPC’s direct target.

*CPC Inhibits Electron Flow through the Electron Transport Chain Starting at Complex I, Nearly as Potently as Rotenone.* To further investigate CPC effects on electron flow through Complex I, via a method fully independent of the TCA cycle, we utilized the MitoCheck Complex I assay. Electrons are fed via NADH to Complex I, and through the full ETC to terminal electron acceptor O_2_ (Fig. 4A). A_340,_ signaling NADH oxidation level (Fig. 4A), is recorded in isolated mitochondria. The assay was employed to measure the effect of 1 h pre-exposure of isolated mitochondria to 0, 10, or 20 uM CPC on electron flow through Complex I. The raw data were processed to calculate rates of change in NADH consumption as detailed in “Materials and Methods.” Fig. 4B shows relative changes in NADH signal due to the various treatments: 0, 10, and 20 uM CPC, or 14 uM rotenone (positive control inhibitor). CPC inhibits the ETC somewhere between Complex I and O_2_, at potency comparable to rotenone, the canonical Complex I inhibitor (Fig. 4C).

*CPC Effects on NADH Oxidation as Low as 10 nM*. Effects of CPC on Complex I of the ETC were further assayed via NADH-Glo luminescent plate-reader assay, which measures the ratio of NAD^+^/NADH in intact cells. RBL-2H3 cells were fed glucose-free galactose-DMEM media, which contains multiple electron sources including amino acids–thus, electrons are flowing from various intake points and through the full ETC to terminal electron acceptor O_2_. Fig. 4D shows low doses of CPC (≥ 0.25 uM; 1 h exposure) inhibit NAD^+^ production by ∼50%. Interestingly, 10 nM CPC consistently stimulated NADH oxidation. However, further investigation of even lower doses (as low as 1 nM) did not show evidence of CPC effect; also apparent inhibition began as low as 50 nM (data not shown).

*CPC Does Not Directly Affect Complex II Activity.* Because electron flow from succinate through cyt C is strongly inhibited by CPC in the Biolog assay (Figs. 2B & 3B), we tested whether CPC directly inhibits Complex II (succinate dehydrogenase) activity, using the MitoCheck Complex II assay. This experimental setup, using isolated mitochondria, measures the flow of electrons from succinate through Complex II, then CoQ (added in excess), and finally to a DCPIP dye which, when reduced, produces signal as A_600_ (Fig. 5A). Concurrently, electron flow from Complex I and to Complex III is shut down by added rotenone and Antimycin A, respectively, isolating effects of Complex II for the assay (Fig. 5A). The assay was employed to measure the effect of 1 h pre-exposure of isolated mitochondria to 0, 10, or 20 uM CPC on electron flow through Complex II. Any potential to observe CoQ inhibition in this assay is masked due to addition of exogenous CoQ, required per manufacturer’s instructions. (However, repeating the assay without using excess CoQ yielded similar experimental results; data not shown).

The raw A_600_ data were processed to calculate rates of electron flow from succinate to the DCPIP dye reduction, as detailed in “Materials and Methods.” Fig. 5B shows relative changes in A_600_ signal due to the various treatments: 0, 10, and 20 uM CPC, or 2.5 mM TTFA (positive control inhibitor). CPC does not inhibit Complex II activity; however, the positive control TTFA did (proving assay integrity), as shown in Fig. 5C. Notably, Complex II is another TCA cycle enzyme (in addition to MDH2) shown to be unaffected by CPC.

*CPC Inhibits Electron Flow through Coenzyme Q, Complex III, to Cytochrome C.* CPC effects on electron flow through Complex III were examined with the use of MitoCheck Complex II/III assay to measure the flow of electrons in isolated mitochondria, from succinate, through Complex II, then CoQ, then Complex III, and finally to cyt C (Fig. 6A). Electron flow was measured via A_550,_ a measure of cyt C reduction. Concurrently, rotenone and KCN are employed to isolate electron flow to Complex II and III (Fig. 6A). Since this assay measures electron flow through both Complex II and III, Complex II inhibition must be (and indeed was, Fig. 5C) eliminated as a potential source of electron flow inhibition before using the Complex II/III assay.

The assay was employed to measure the effect of 1 h pre-exposure of isolated mitochondria exposed to 0, 10, or 20 uM CPC on electron flow between Complex III and cyt C in isolated mitochondria. The raw data were processed to calculate rates of cyt C reduction. Fig. 6B shows relative changes in cyt C signal (A_550_) due to the various treatments: 0, 10, and 20 uM CPC, or 28 uM Antimycin A (positive control inhibitor).

CPC significantly inhibits electron flow from succinate, at some point at or after CoQ (Fig. 6C). *CPC Inhibits Electron Flow from Cytochrome C to Complex IV, as Potently as Cyanide.* To investigate CPC effects on electron flow through Complex IV, we utilized MitoCheck Complex IV assay to measure the flow of electrons, in isolated mitochondria, from cyt C to Complex IV and terminal electron acceptor O_2_ (Fig. 7A). Electron flow was measured via A_550,_ an indicator of cyt C redox status. Potassium cyanide (KCN), the positive control, requires use at elevated pH (7.8) for sufficient deprotonation and thus activity (detailed in “Materials and Methods”). Therefore, CPC and KCN were tested in separate buffers (Fig. 7C): CPC in “Control_CPC_” MitoCheck Complex IV buffer at pH 7.4, and KCN in “Control_KCN_” buffer at pH 7.8 (the same buffer plus added 0.05 M NaOH). For the other set of experiments (Fig. 7D), all samples tested (CPC or KCN) were dissolved in the same 0.05 M NaOH-containing Complex IV buffer, for direct comparison of CPC with KCN (this is simply called “Control” in Fig. 7D).

The assay was employed to measure the effect of 1 h pre-exposure of isolated mitochondria to 0, 10, or 20 uM CPC; 10 uM KCN; or 10 or 20 uM hexadecylbenzene (HDB) on electron flow between cyt C and Complex IV. The raw data were processed to calculate rates of cyt C oxidation, as detailed in “Materials and Methods.” Fig. 7B shows relative changes in cyt C signal due to the various treatments.

CPC inhibits electron flow between cyt C and Complex IV as potently as KCN in both individualized (Fig. 7C) and standardized (Fig. 7D) buffer conditions. However, HDB, an analog of CPC which lacks its positively-charged head group, has no effect on electron flow in this assay (Fig. 7E).

*CPC Boosts Reduction of CoQ.* Effects of CPC on CoQ redox state were assayed via Arigo colorimetric dye (A_620_) which absorbs upon interaction with CoQ’s hydroxyl group when it is reduced. Therefore, the assay detects the reduction state of and, therefore, electron flow to CoQ. The assay was employed to measure the effect of 1 h pre-exposure of intact RBL-2H3 cells to 0, 5, or 10 uM CPC on electron flow to CoQ in BT-glucose media. The raw data were processed to calculate the amount of reduced Q, normalized to the control, as detailed in the “Materials and Methods.” Fig. 8 shows CPC causes accumulation of reduced CoQ in intact cells.

**Figure 8.**
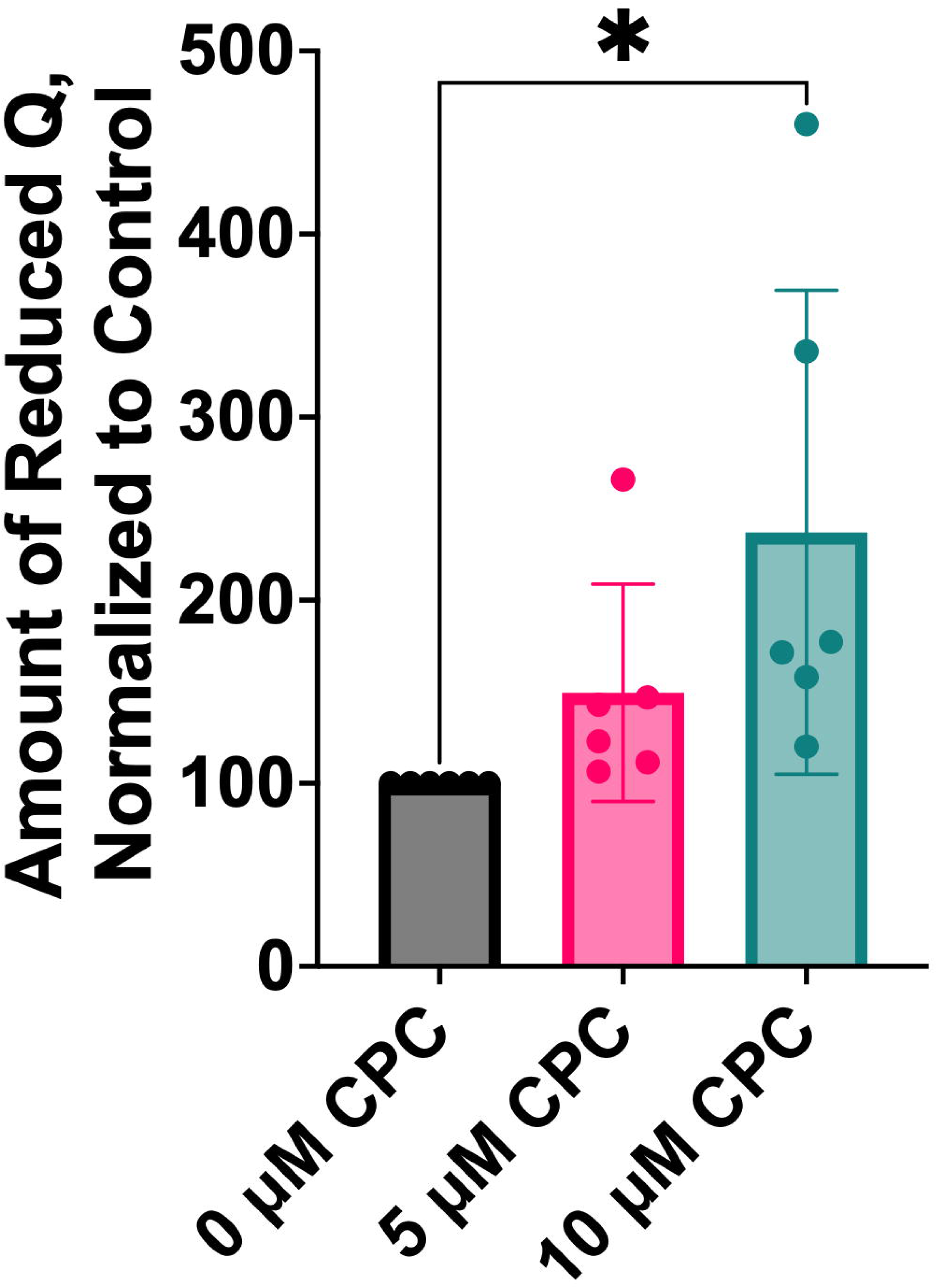
Effect of CPC on electron flow to CoQ in intact RBL-2H3 cells. Arigo Q reduction assay was used to measure the concentration of reduced CoQ (read via A_620_, plate reader) in RBL-2H3 cells. For each experiment, intact cells were pre-exposed to CPC for 1 h in glucose-BT, then lysed via sonication. Lysates were analyzed via Bradford protein assay to standardize signals. CoQ levels were normalized to the no-treatment control of each given day and are shown as mean ± SEM from 6 days of experiments (duplicates per experiment). Significance analysis was performed via one-way ANOVA with Dunnett’s post-hoc test, *p < 0.05.

The assay was repeated in glucose-free galactose-DMEM to determine whether forcing cells to use their mitochondria for energy production would affect results. The trend of CoQ reduction was the same regardless of buffer change (data not shown).

*CPC Does Not Alter Total Cellular Levels of Mitochondrial Lipid Cardiolipin.* To characterize acute CPC effects on cardiolipin (CL) levels, a fluorescent probe from Abcam CL assay was employed, as described in “Materials and Methods.” This assay was used to measure the effect of pre-exposing RBL-2H3 cells in BT-glucose for 1 h to 5 or 10 μM CPC. Under these conditions, CPC does not affect total levels of CL (Fig. 9).

**Figure 9.**
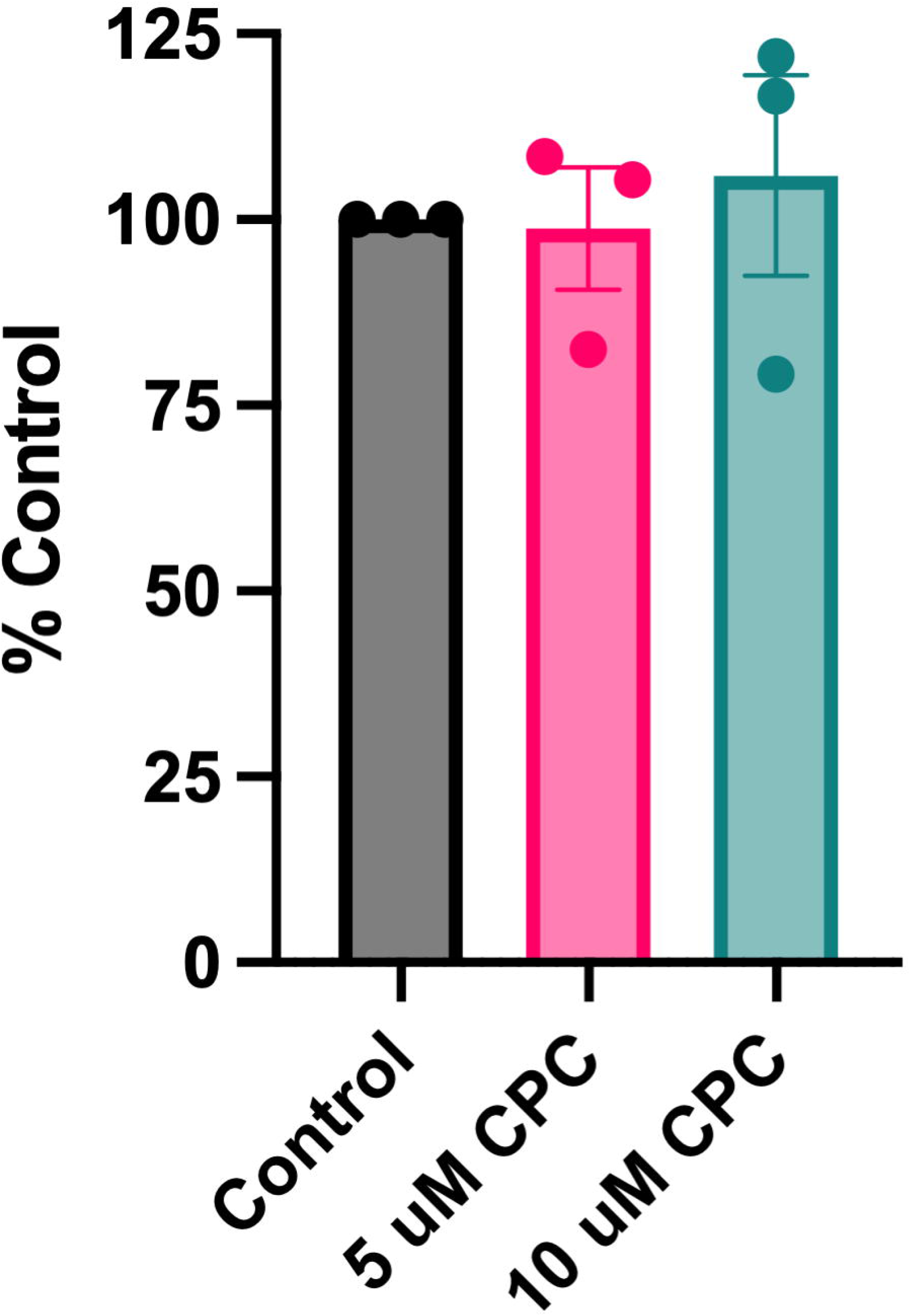
Total cardiolipin levels in RBL-2H3 cells. Intact RBL-2H3 cells were exposed to CPC for 1 h in glucose-BT, then lysed via sonication to determine cardiolipin levels via fluorescent dye. Lysates were analyzed via Bradford protein assay to standardize signals. Cardiolipin levels were normalized to the no-treatment control of each given day and are shown as mean ± SEM from 3 days of experiments (duplicates per experiment). Significance analysis was performed via one-way ANOVA with Dunnett’s post-hoc test.

The assay was repeated in glucose-free galactose-DMEM to determine whether forcing cells to use their mitochondria for energy production would affect results. Also in this media, CPC does not alter total cellular CL levels (Fig. S7).

*CPC Interferes with the Binding of Mitochondrial Lipid Cardiolipin to its Detection Probe*. CPC effects on the ability of purified CL to bind its fluorescent probe (Abcam) were investigated *in vitro*. First, CL (0, 1, and 10 uM) was exposed in assay buffer (Abcam) to CPC (0, 5, or 50 uM) for 15 min. Next, Abcam’s proprietary CL-binding probe was added to the solutions. The dye fluoresces when it binds to CL. At the CPC doses tested, dye fluorescence declined significantly, at both CL doses assessed (Fig 10A).

**Figure 10.**
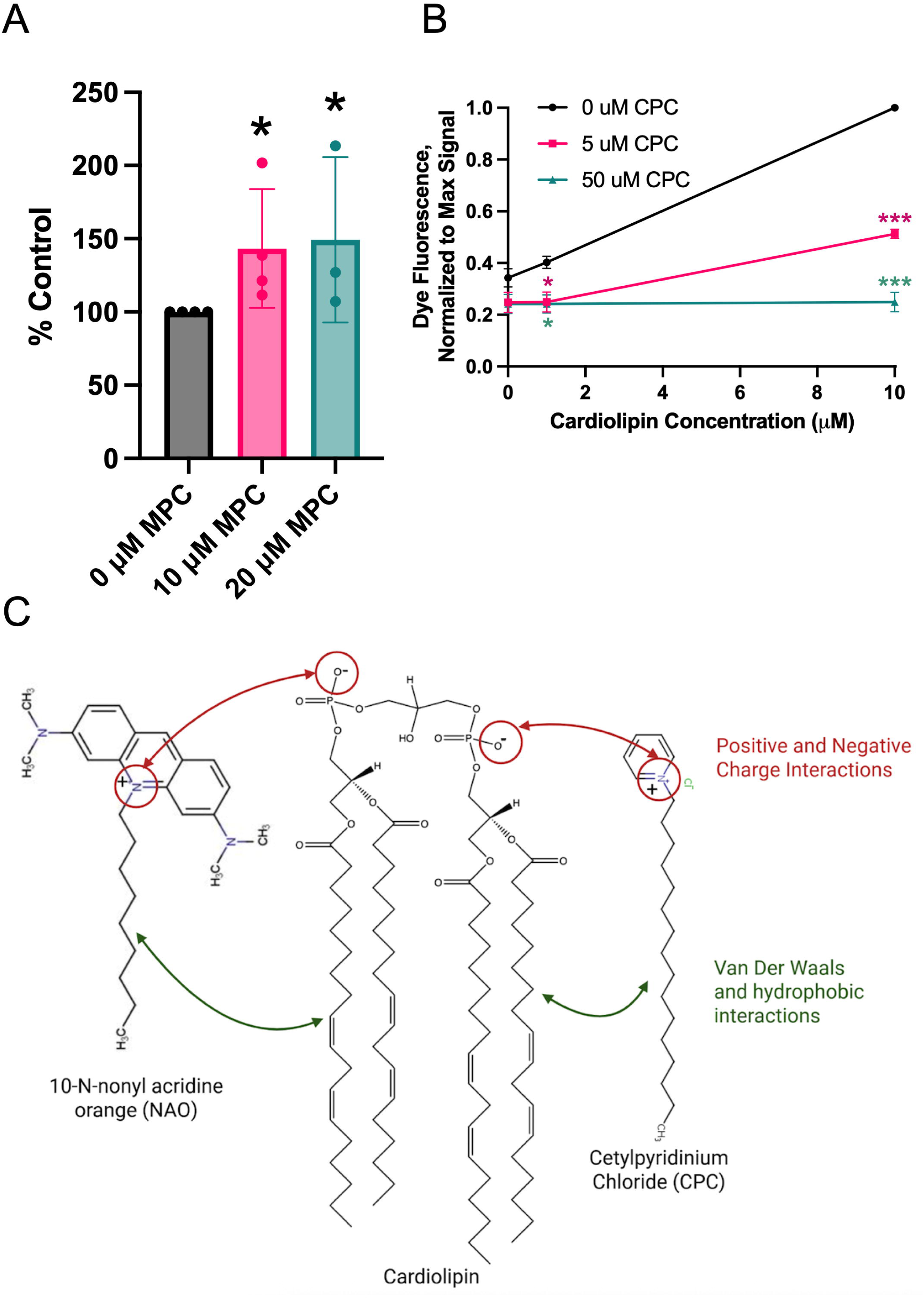
Assessment of CPC interference with cardiolipin binding to its partners. (**A**) Cardiolipin was pre-exposed to CPC (at the indicated doses) for 15 min in glucose-BT buffer then was detected (fluorescence, via plate reader) by Abcam’s fluorescent cardiolipin probe (used in Fig. 9). Probe fluorescence was normalized to maximum signal (10 μM cardiolipin, without CPC) and plotted as a function of cardiolipin present. Data were gathered from 3 experiments. Significance was determined by one-way ANOVA with Dunnett’s post-test comparing CPC treatment to control at each concentration of cardiolipin, ***p < 0.001, *p < 0.05. (**B**) Cardiolipin-coated beads (Echelon) were pre-exposed to CPC’s head group (MPC) for 1 h, then incubated for 3 h with cytochrome c ± MPC. Cytochrome C levels in supernatant (unbound) were measured by A_410_ and presented as % of untreated control. Data were gathered from 3-4 experiments. Significance was determined by one-tailed, nonparametric t-test compared to control value, *p < 0.05. Data in each graph are displayed as mean ± SEM. (**C**) On the left is shown the mechanism of binding between cardiolipin and a specific probe. On the right, a proposed mechanism for CPC’s noncovalent interactions with cardiolipin. Created in Biorender.

*Analog for the Head Group of CPC Interferes with Binding of Cardiolipin-Coated Beads to Cytochrome C*. To determine whether CPC affects the binding of CL to its critical protein partner cyt C, a novel assay was developed, using CL-coated beads and purified cyt C. MPC (1-methylpyridinium chloride) was used to represent the positively-charged nitrogenous head group of CPC, with a methyl group in place of CPC’s lipid tail. Following 1 h pre-exposure of the CL-coated beads to 0, 10, or 20 uM MPC, cyt C was added to the bead mixtures with fresh MPC for a 3 h incubation. Next, A_410_, measuring total cyt C, was used to probe the amount of cyt C remaining unbound to CL. MPC significantly increased the amount of unbound cyt C (Fig. 10B). Thus, MPC, CPC’s head group, blocks the binding of CL to its critical mitochondrial binding partner cyt C.

## Discussion

CPC is used widely in over-the-counter products and food, yet researchers have not yet determined exposure levels that may harm health via its mitotoxic (Weller et al., Datta, Chavez, Saladino) outcomes. Toxicological science supplies consumers and regulators with data to make informed decisions. While the U.S. FDA has effectively banned CPC from hand soaps and sanitizers (20, 21), CPC remains in many products, largely unregulated. For comparison, another widespread compound, triclosan, was used in products for decades until several studies revealed toxic effects^43,28,42-45^. Despite the many links between mitochondrial toxicity, like effects caused by CPC(Chávez & Bravo, 1982a; Datta, He, et al., 2017b; Liu et al., 2023; Saladino et al., 1971; Weller et al., 2024) and human disease(Arron et al., 2024; Bhandari et al., 2014; Cataldo et al., 2010; Cui et al., 2010; Hara et al., 2014; Jheng et al., 2012; Pozo Devoto & Falzone, 2017; Ryter et al., 2018; Tomas et al., 2017; Youle & Van Der Bliek, 2012), researchers have not yet connected CPC exposure to health effects via epidemiology (Pubmed, 2026). Knowledge of biochemical mechanisms underlying CPC’s mitotoxicity is essential to focus future epidemiological investigations onto relevant diseases. Mechanistic data also presents opportunities for predictive toxicology, including the vast class of quat compounds.

In this study, we aimed to determine the direct target of CPC responsible for its inhibition of OCR and, thus, mitochondrial function. Previous researchers did not investigate CPC effects on the full ETC–there is a lack of information on CPC’s interactions with Complexes III and IV, coenzyme Q, or cytochrome c.

Biolog MitoPlate S-1 electron flow data (Figs. 2 & 3) display apparent CPC disruption of Complex I & II as well as of CoQ. These experiments monitored electron flow from several ETC-feeding substrates, in both primary human keratinocytes and RBL-2H3 Cells. For many substrates, effects were potent even at acute exposure with 0.5 uM. CPC inhibits electron flow from Complex I-feeding substrates (Fig 2A, 3A) malate, fumarate, α-ketoglutarate, and Ala-Gln. (The lack of significance of the fumarate data in the RBLs may be due to the relatively low A_590_ signal from this substrate in RBLs, versus in the keratinocytes, which did produce robust electron flow from fumarate [Table S2] and evinced CPC inhibition.) Also, CPC inhibits electron flow from Complex II substrate succinate (Fig 2B, 3B) and CoQ-feeding substrate α-glycerol phosphate (Fig. 2C, Fig 3C). However, the Complex I, Complex II, and CoQ-feeding substrates could be apparently inhibited due to downstream effects since the assay reads out after cyt C (Fig. 1). (Note, additional TCA substrates, such as citrate, are included on the MitoPlate S-1, but there was insufficient electron flow signal from these in controls, under the conditions tested.)

The Biolog results show that CPC inhibits electron flow at some point between the individual substrates fed to the cells and cyt C, which is directly before detection via redox dye (Fig 1). Many of the substrates require processing by TCA enzymes before they can feed electrons into the ETC. Thus, to test CPC effects on the TCA cycle itself, we focused on a TCA substrate that produced robust effects in Biolog (Figs. 2 & 3) and developed an enzyme assay for TCA enzyme mitochondrial malate dehydrogenase (MDH2). CPC did not affect MDH2 activity (Fig. S2). Additionally, succinate dehydrogenase (Complex II), a second TCA enzyme, was also tested and found to be unaffected by CPC (Fig. 5, to be discussed below). Thus, two of the eight TCA enzymes, representing disparate portions of the cycle, are unaffected by CPC. Furthermore, Mitocheck experiments cut out the TCA cycle altogether (in contrast to the Biolog experiments), yet CPC still affects electron flow in several of these isolated mitochondria assays, further moving the focus away from the TCA cycle.

Therefore, to focus on the ETC and to further probe Complex I, we moved to MitoCheck assays in isolated mitochondria (Fig. 4 A-C) as well as NADH-Glo experiments in intact cells (Fig. 4D). We found that CPC inhibits electron flow in isolated mitochondria via Complex I with comparable potency to that of canonical Complex I inhibitor rotenone (Fig. 4C). The Mitocheck assay for Complex I of the ETC was used to detect CPC inhibition of electron flow from NADH through Complex I–*including* through the remainder of the ETC (Fig. 4A). Also, NADH oxidation in intact RBL-2H3 cells is inhibited by CPC (Fig 4D), as low as 0.25 uM. Like the Biolog assay (Figs. 2A & 3A), the NADH-Glo assay shows a 50% inhibition by ≤ 1 uM CPC (Fig 4D). Thus, these three disparate assays all indicate CPC-disrupted electron flow from NADH, through Complex I (Fig. 2A, 3A, 4C, 4D), then farther along the ETC, either to cyt C (Fig. 1) or all the way to O_2_ (Fig. 4A). Each of these three assays allow electrons to continue flowing through the ETC, far past Complex I. Thus, Complex I is not necessarily the specific site of inhibition; it is equally likely the inhibition of NADH oxidation occurs due to downstream interference within the ETC.

To train focus on Complex II next, we performed a MitoCheck assay in isolated mitochondria (Fig. 5). The Mitocheck assay for Complex II (succinate dehydrogenase) was used to detect CPC effects on electron flow from succinate through Complex II, CoQ, and to the DCPIP dye (Fig. 5A). In contrast to the Biolog findings (Fig. 2B & 3B), CPC does not impact electron flow in this assay of isolated mitochondria through Complex II (Fig. 5B & 5C), even as positive control TTFA was inhibitory (Fig. 5C). This contradiction is explained by the setup of these methods: Complex II Mitocheck assay isolates electron flow from succinate through CoQ, to the DCPIP dye (Fig. 5A)--whereas the Biolog assay measures electron flow from succinate all the way through cyt C. Thus, in combination, these two assays point toward a CPC target *after* CoQ of the ETC.

Of note, the Complex II Mitocheck protocol calls for addition of excess CoQ, likely to enhance signal level, but which could mask potential CPC inhibition of native CoQ. However, repeating the assay without using excess CoQ yielded similar experimental results, as noted in “Results” –suggesting that CPC does not inhibit CoQ. Moving further down the ETC, the MitoCheck Complex III assay functions by feeding succinate to isolated mitochondria and measuring cyt C reduction (Fig. 6A). Therefore, CPC interference with Complex II, CoQ, Complex III, *or* cyt C could produce the same result (Fig. 6C). Recall that the Complex II MitoCheck assay requires the addition of exogenous CoQ; however, the Complex III MitoCheck assay does not. Thus, any direct inhibition/stimulation of endogenous CoQ function by CPC is measured. CPC is inhibitory in this assay, as is positive control Antimycin A (Fig. 6C).

Interestingly, apparent Complex I inhibition (Fig. 4C) is similar in magnitude to that of Complex III (Fig. 6C). Since the Mitocheck Complex I assay feeds electrons through both Complex I and III, it is likely that if both were directly inhibited, we would see stronger inhibition for the Complex I assay as compared to the Complex III assay. These findings further implicate a CPC effect late in the ETC, after Complex III. Additionally, comparison of the experimental setups of MitoChecks for Complex I (Fig. 4A) and for Complex III (Fig. 6A) shows that I measures through to O_2_ whereas III stops at cyt C. The fact that both are nearly equally inhibited rules out Complex IV as the direct target of CPC and points a finger in the direction of cyt C.

However, we proceeded next with the MitoCheck Complex IV assay in isolated mitochondria, to gain further mechanistic insight. Fig. 7A displays the experimental schematic, in which reduced cyt C is fed to mitochondria and direct assessment is made of its oxidation by Complex IV through to O_2_. The apparent decrease in Complex IV activity is the most dramatic of all the Complexes tested by Mitocheck assays (Fig. 7B-7D). To test whether the cause was electrostatic in nature, HDB (CPC without the positive head-group) was used. HDB did not affect Complex IV activity (Fig. 7E), indicating the positively-charged nitrogen of CPC and, therefore, likely electrostatic interference, is required to induce the inhibition. We also carefully compared CPC to cyanide, the canonical Complex IV inhibitor, and found that CPC is as potent as cyanide (via a separate ANOVA with Tukey’s post-test comparing 10 uM CPC to 10 uM cyanide and indicating lack of significant difference between CPC and cyanide). However, the analysis of Complex I vs III data noted above strongly signals that CPC does not directly affect Complex IV. This combination of results leads to the conclusion that CPC, instead, targets the other player in the Complex IV assay: cytochrome C.

Yet, the reason CPC decreases activity of Complex IV more than that of Complexes I & III (assayed in a manner also employing cyt C) remains unknown. Perhaps attempted adaptation actually enhances electron flow through earlier parts of the chain, which are not active in the assay for Complex IV (Fig. 7A, due to the only electron source in the Complex IV assay being, reduced cyt C).

In fact, CPC stimulates reduction of CoQ in intact cells fed glucose (Fig. 8), a situation in which electrons are incoming to the ETC via various paths (including Complex I and II). The observed increase in reduced CoQ shows that CPC’s target to inhibit electron flow is not upstream of CoQ–CPC’s target is *not* Complex I, Complex II, or α-glycerol-3-phosphate dehydrogenase (Fig. 1), nor the TCA cycle. The buildup of reduced CoQ, however, without subsequent oxidation of CoQ by Complex III, further implicates CPC inhibition to be occurring proximally yet downstream of CoQ, including but not isolated to our theory that CPC acts via cytochrome C disruption (Figs. 10C and 11). This increase in reduced CoQ also indicates Complex I, which feeds into CoQ, is itself not inhibited, and the apparent Complex I inhibition (Fig. 4C) is due to downstream blockage. Additionally, α−ketoglutarate feeds electrons to CoQ (Fig. 1). However, in the Biolog assay where electron flow from α-glycerol-3-phosphate appears to be blocked by CPC (Figs. 2C & 3C), the electron path continues downstream, through to cyt C (also as in the Complex I assays). The increase in reduced CoQ also indicates CoQ isn’t being oxidized by Complex III, and thus there is a blockage closely after or at Complex III, stopping electrons from continuing from CoQ through the ETC. The Biolog ETC assay only measures electron flow through cyt C, ending in the reduction of MC redox dye (after cyt C). This would suggest the inhibition is after CoQ and before Complex IV, confirming all previous results thus far.

After CoQ, electrons are fed to Complex III, and then to cyt C. All CPC effects have now been narrowed down to this point in the ETC: after Complex III. Cyt C is attracted to the IMM largely through noncovalent interactions between cyt C’s cationic residues (lysine) and anionic phospholipids on the outer leaflet of the IMM, mainly cardiolipin (Fig. 10C)(Osheroff et al., 1983; Rytömaa et al., 1992; Rytömaa & Kinnunen, 1994; Vik et al., 1981). This cyt C-cardiolipin interaction is necessary to draw the aqueous protein cyt C out of the intermembrane space and toward the IMM, in which its partners Complexes III and IV reside.

In fact, cyt C-cardiolipin is further implicated as the target of CPC in the mitochondria because of the lack of Complex II inhibition by CPC. The Complex II MitoCheck assay is the only one of the four which does not include cyt C (compare schematics in Fig. 5A to Figs 4A, 6A, and 7A)--and it is the only one unaffected by CPC. Thus, we tested whether the short-term CPC exposures used in this study affect gross levels of cardiolipin (CL) in RBL cells. To do so, a proprietary fluorescent CL probe was employed. This experiment was done both in glucose (Fig. 9) and in glucose-free galactose-DMEM (Fig. S8). These data reveal that total cellular levels of CL are unaffected by CPC. Therefore, the mitochondrial effects described in this paper are likely not due to gross alteration of CL levels. However, it remains possible that chronic CPC exposure could alter CL levels because CPC blocks the lipid phosphatidylinositol 4,5-bisphosphate (PIP_2_)(Raut, Weller, et al., 2022), which is needed to synthesize CL(Paradies et al., 2019) via CL synthase (which uses diacylglycerol made from PIP_2_). CPC’s perturbation of PIP_2_ could manifest as a decrease in CL levels during long-term exposures but would go unseen in acute exposure conditions. Disruption of CL production is associated with the grave genetic disorder due to enzyme mutation, Barth syndrome(Gebert et al., 2009). CL dysfunction in Barth’s syndrome is likely responsible for the defects to mitochondrial structure and function seen in cardiac muscle and neutrophil bone marrow cells.

In the absence of gross changes in CL quantity, we turned toward testing direct CPC chemical effects on CL. In an *in vitro* test, we examined CPC effects on the binding of CL to the same proprietary fluorescent dye used in Fig. 9. While the company would not reveal the identity of this dye, it maybe similar to nonylacridine orange (NAO)(Petit et al., 1992) or TTAPE-Me (Petit et al., 1992), compounds which specifically bind and detect CL via an electrostatic and possibly also hydrophobic mechanism(Leung et al., 2014). Interestingly, like CPC, CL-binding dyes NAO and TTAPE-Me are quats: potential electrostatic interaction between NAO, CPC, and CL is depicted in Fig. 10C. The probe’s fluorescence is enhanced in the presence of increasing CL concentrations (black line in Fig. 10A). The CL probe’s fluorescence was indeed diminished by CPC exposure (Fig 10A), suggesting that CPC binds CL, displacing the probe. Thus, we hypothesized that CPC may similarly bind CL within mitochondria in a manner that would displace cytochrome C from its critical lipid binding partner.

To test this hypothesis, CL-coated beads were used to conduct another CL binding assay, this time between the CL on the beads and purified cyt C (Fig. 10B). In the control (no chemical treatment, “0 uM MPC” in Fig. 10B), some of the added cyt C was in excess and remained in the supernatant following 3 h of incubation of the protein with the CL beads. Treated samples utilized MPC as a proxy for CPC (MPC is CPC with a methyl group in place of CPC’s 16-carbon lipid tail; CPC structure in Fig. 10C).

The reason for using MPC in place of CPC is that this Echelon lipid bead assay requires the use of the detergent Tween-20 (for reduction of nonspecific binding), which causes CPC to be bound up into micelles, reducing its availability to interact with biomolecules (Ghosh, 2001) in such a way that might reduce its ability to directly bind lipids like CL. To avoid this complication, we instead employed MPC. Additionally, we utilized a low (10 mM NaCl) salt concentration similar to that found in the cytosol and intermembrane space, for physiological relevance because NaCl concentrations are much lower in the cytosol (and thus the intermembrane space), compared to the blood/extracellular space(Baggaley & Riedel, 1966). Use of low salt is also to avoid the experimental complication that binding of CL to positively-charged partners is known to be interrupted by high salt(Hanske et al., 2012) due to Debye screening, making interactions weaker than they are in living mitochondria. Under these carefully designed conditions, we found that MPC causes cyt C to fall off the CL-coated beads (Fig. 10B).

Together, these CL binding experiments (Fig. 10A & 10B), along with the experiments in Figs. 2-9, suggest that direct interference (Fig. 10C) of CL is CPC’s likely sole target within the ETC (Fig. 11). Considering the positively-charged nitrogen of CPC and negatively charged phosphate groups of CL, in comparison to the electrostatic interaction between the quat dyes (TTAPE-Me and NOA) and CL, we propose the aforementioned CPC interference is likewise via electrostatic interference. This CL hypothesis is bolstered by the fact that CPC is already well-documented to interfere with another critical signaling lipid, PIP_2_, in immune cell signaling events(Raut, Weller, et al., 2022). PIP_2_ is highly similar in structure to CL. Both lipids contain hydrocarbon chains embedded in the lipid bilayer, a glycerol backbone, and multiple negatively charged phosphates exposed to the aqueous side. CPC directly disrupts the binding of PIP_2_ to several of PIP_2_’s binding partners: 1.) influenza protein hemagglutinin’s basic residues (arginine and lysine)(Raut, Weller, et al., 2022), 2.) influenza protein M1(Raut, Obeng, et al., 2022), 3.) immune signaling protein MARCKS in MCs(Raut, Weller, et al., 2022), 4.) Pleckstrin Homology reporter constructs(Raut, Obeng, et al., 2022), and 5.) the Spike protein of SARS-CoV-2(Raut, Waters, et al., 2022b). Additionally, CPC disrupts clustering and dynamics of PIP_2_ at the plasma membrane(Aho et al., 2025). Given these known effects on PIP_2_ along with the experimental results in Fig. 10, CPC may similarly disrupt cyt C binding to cardiolipin.

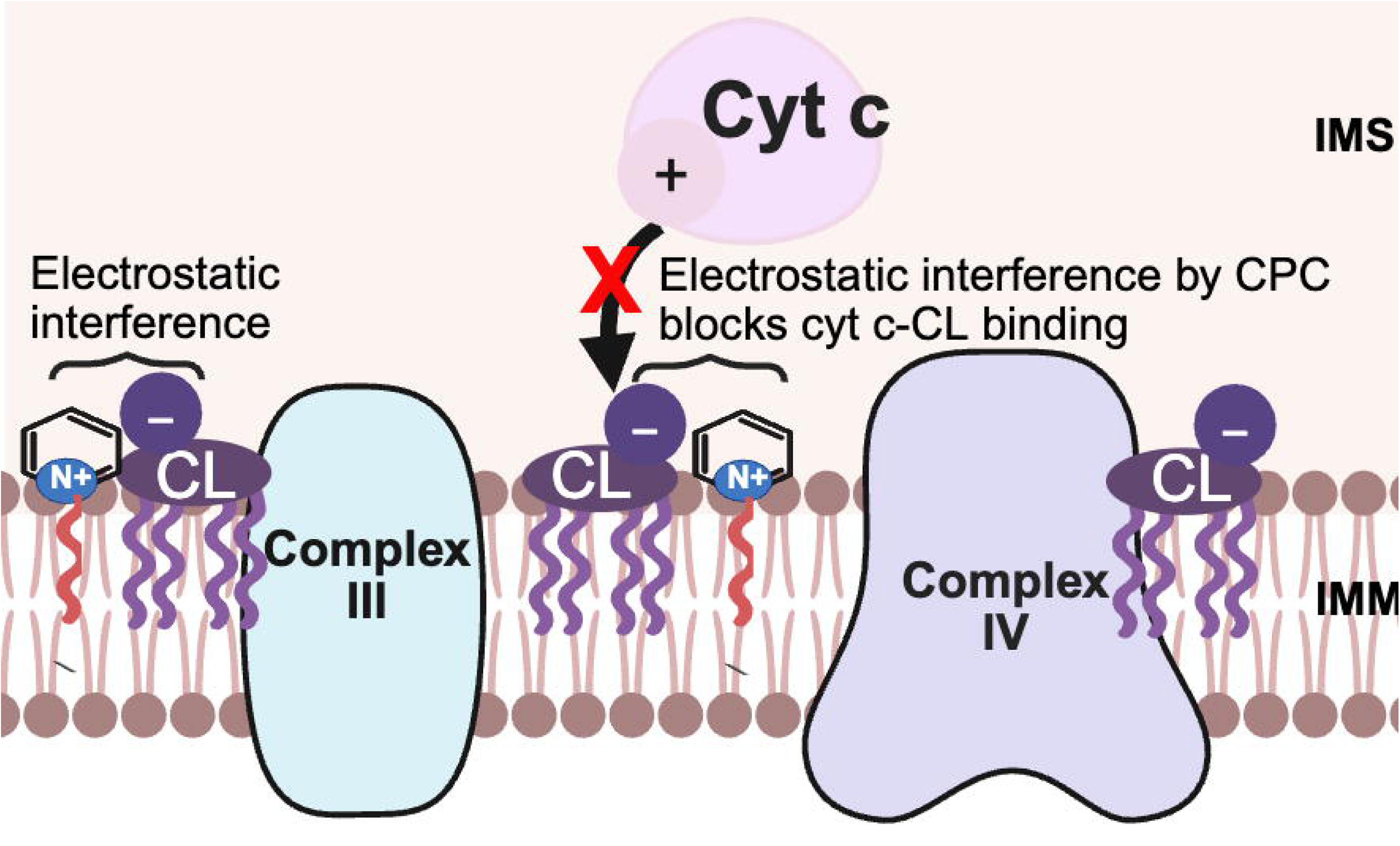

It is important to note that not all lipids are disturbed by CPC: the activity of critical electron transfer lipid CoQ is uninhibited by CPC–CoQ actually collects more electrons during CPC exposure. Additionally, the plasma membranes of cells treated with this study’s exposure conditions were unaffected in structure and membrane potential(Obeng et al., 2023).

CPC is not the first toxicant to be implicated in disruption of CL-cyt C binding, as an underlying mechanism of mitochondrial toxicity. Positively-charged cadmium was also shown to do so, via an electrostatic mechanism(Romanova et al., 2025). Additionally, electrostatic CL interactions with -synuclein are implicated in Parkinson’s disease (reviewed in (Ruiz-Ortega et al., 2025), shining a light on the question of whether CPC could similarly affect Parkinson’s risk. In general, perturbations in CL are associated with neurodegeneration (Ruiz-Ortega et al., 2025).

In an unexpected turn, we found CPC to be a more potent mitochondrial toxicant in cells versus in isolated mitochondria, despite the same exposure times and despite the reputation that isolated mitochondria have as being more fragile and susceptible8/18/2026 12:47:00 PM. In cells, including in primary human keratinocytes, we found inhibitory effects as low as 0.25 uM (Fig. 4D). In contrast, the experiments using isolated mitochondria (Figs. 4, 5, 6, and 7) sometimes required 10 uM for observing CPC effects. This disparity may reflect the ability of cells with even semi-intact plasma membranes to concentrate CPC, allowing for effects at much lower doses when cells are used. This comparative effect implies that CPC is actually more potent in real-life biological exposures than in isolated mitochondrial experiments.

Notably, the Mitocheck experiments employ as positive controls known canonical mitotoxicants with specific known mechanisms on individual respiratory Complexes. Complex I results compare the effect of 14 uM rotenone to 10 or 20 uM CPC, revealing a similar inhibitory effect of CPC as has this causative agent of Parkinson’s(Dhillon et al., 2008). While CPC at 20 uM was less potent than 28 uM Antimycin A on Complex III (Fig. 6), both chemicals were inhibitory. Finally, in Mitocheck Complex IV, canonical Complex IV-inhibitor KCN was used at 10 uM and affected electron flow through Complex IV on par with 10 uM CPC. Although, as described in “Materials and Methods,” KCN is truly added as 250 uM, while 10 uM is the estimated active concentration due to pH/protonation–whereas CPC is truly delivered at the indicated doses.

Interestingly, another widespread quat, benzalkonium chloride (BAK) has been carefully investigated to determine its mechanism of mitotoxicity. The researchers found that BAK specifically targets *only* Complex I (Datta, Baudouin, et al., 2017b). Their systematic experiments and multiple approaches working through the ETC showed that, in stark contrast to CPC, BAK does not inhibit CoQ, cyt C, or Complexes III or IV. BAK’s structure is very similar to that of CPC, apart from its shorter hydrophobic chain length ahead of the quaternary ammonium. This contrast suggests that CPC’s hydrophobic chain of 16 carbons is critical to its affinity for cardiolipin and thus toxicity toward Complexes III and IV and cyt C.

In summary, here we have a revealed novel mechanism of mitochondrial toxicity by the common-use antimicrobial CPC on the ETC. We show that CPC potently inhibits electron flow in primary human keratinocytes, RBL cells, and isolated mitochondria. Considering that together all evidence presented in this paper, we find that the culprit behind the ETC chaos is electrostatic CPC interference with the anionic lipid cardiolipin and its critical binding partner, cationic cyt C (Fig 11). This CPC interference of CL-cyt C binding explains the apparent inhibition of all ETC components, as noted in Table 1. However, closer investigation showed that Complexes I and II and CoQ are actually not harmed directly by CPC (Table 1). Activities of Complexes III and IV are blocked by CPC albeit indirectly (Table 1). The only remaining core ETC component to be pinpointed by CPC is cytochrome C and its lipid partner CL. This work demonstrates a novel mechanism of toxicity (Fig 11) for CPC and has implications for both epidemiology of CPC and predictive toxicology generally. The work reveals novel mechanisms by which CPC may impact human health.

**Table 1.** Summarizes the electron flow inhibition findings of this manuscript.

| ETC Target | Inhibited by CPC? | Indirect, Direct, or Not Inhibited? |
| --- | --- | --- |
| Complex I | Apparent Inhibition | Likely Not Inhibited |
| Complex II | Apparent Inhibition | Not Inhibited |
| α-Glycerol-3-phosphate Dehydrogenase | Apparent Inhibition | Likely Not Inhibited |
| Coenzyme Q | Apparent Inhibition | Not inhibited |
| Complex III | Apparent Inhibition | Indirect |
| **Cytochrome c/ Cardiolipin** | **Yes** | **Direct** |
| Complex IV | Apparent Inhibition | Indirect |

## Acknowledgements

We thank Dr. Bright Obeng for cell maintenance and lab support; Tania Systuk, Sydni Plummer, Isaac Dostie, and Jeongwon Eom for lab support; Sasha Weller for idea generation; Dr. Jennifer Newell-Caito for insightful discussions, and Dr. Joshua Kelley for allowing our lab to use their Sonifier.

## Supplementary material

Supplementary material is available online.

