## Supplement for "Antimicrobial cetylpyridinium chloride disrupts the mitochondrial electron transport chain as potently as cyanide, via cardiolipin interference at cytochrome C"

**Table of Contents**

**Fig. S1. Effect of CPC on Biolog MC Redox Dye** …………..………………………............………...3

**Table S1. Substrates Statistically Selected for Further CPC-Effect Analysis**……...…….………4

**Table S2. Substrate Slope Parameters for Biolog Electron Flow Rate Assay Analysis** ……….5

**Fig. S2. Effect of CPC on TCA cycle Enzyme Mitochondrial Malate Dehydrogenase (MDH2) Enzyme Activity**…….……………………………………………………………………………….……….6

**Table S3. Readout Absorbances Used for Mitocheck Assays**………………………………………8

**Fig. S3. Justification for Mitocheck Complex I Conditions and Analysis Parameters: Time Range Selection, Absorbance Range Measured, and Vehicle Buffer** **Effects** …...…………………………………………………………………………………..…………………….……9

**Fig. S4. Justification for Mitocheck Complex II Conditions and Analysis Parameters: Time Range Selection, Absorbance Range Measured, and Vehicle Buffer** **Effects**…...…………………………………………………………………………………..……………….10

**Fig. S5. Justification for Mitocheck Complex III Conditions and Analysis Parameters: Time Range Selection, Absorbance Range Measured, and Vehicle Buffer** **Effects**……………………………………………………………………………………………..………….11

**Fig. S6. Justification for Mitocheck Complex IV Conditions and Analysis Parameters: Time Range Selection, Absorbance Range Measured, and Vehicle Buffer** **Effects**………..………….12

**Fig. S7. Effect of CPC on Cardiolipin Levels in Glucose-free Galactose-DMEM**………………. 13

**Fig. S1. Effect of CPC on Biolog MC Redox Dye**

**Method:** In order to determine whether CPC interferes with the Biolog MC Redox Dye, an analysis of Biolog Electron Flow Assay data from in RBL-2H3 mast cells was conducted. To assess the CPC effect on MC redox dye, the maximum and minimum dye absorbance signal resulting from the Biolog Electron Flow assay (see “Materials and Methods”) in each no-substrate control well, were analyzed as a function of 0 uM, 0.5 uM, and 1.0 uM CPC exposure. A small electron flow rate was, in fact, registered even in unfed no-substrate wells, likely due to leftover electron sources even in washed cells. This no-substrate A_590_ electron flow value is roughly 1 to 20-fold lower than that produced by added test substrates (compare Fig. S1 y-axis values to Table S2 A_590_ values).

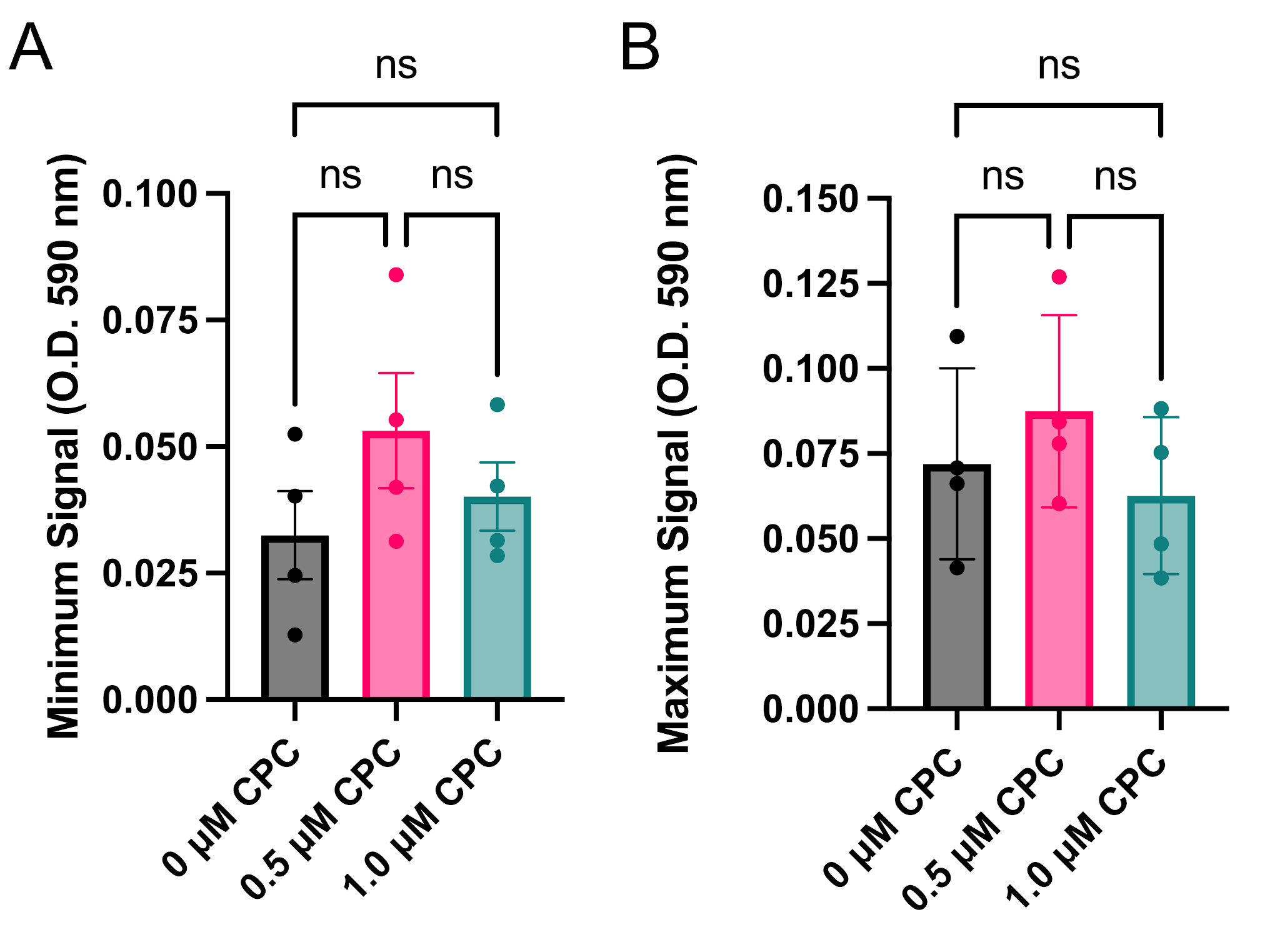

**Figure S1.** **Control assessing whether CPC interferes with the Biolog MC redox dye**. Electron flow was measured in saponin-permeabilized RBL-2H3 cells (via A_590_, plate reader) from no-substrate control samples, through cyt C, during 3 h exposure to various CPC concentrations in MAS buffer. The y-axis represents the rate of electron flow in unfed cells. **(A)** Effect of 0, 0.5, and 1.0 uM CPC on minimum A590 signal (over the course of the 3 h), for each no-substrate control well. **(B)** Effect of 0, 0.5, and 1.0 uM CPC on maximum A_590_ signal (over the course of the 3 h), for each no-substrate control well. Data shown are means ± SEM for 4 experiments. Significance was tested via one-way ANOVA with Tukey’s post-hoc, ns = not significant.

**Result:** CPC does not affect the Biolog MC redox dye absorbance signal.

**Conclusion**: Any effects due to CPC in Biolog electron flow data are due to physiological effects of CPC on the cells, rather than artifacts due to dye interference.

**Table S1. Substrates Statistically Selected for Further CPC-Effect Analysis**

|  | **P-Value** | |
| --- | --- | --- |
| ***Substrate*** | **Primary Human Keratinocytes** | **RBL-2H3** |
| ***malate*** | **0.009** | **0.008** |
| ***⍺-glycerol phosphate*** | **0.058** | **0.058** |
| ***⍺-ketoglutarate*** | **0.009** | **0.042** |
| ***fumarate*** | **0.06** | **0.032** |
| ***succinate*** | **0.014** | **0.012** |
| ***Ala-Gln*** | **0.100** | **0.084** |

**Methods**: In order to select for analysis the substrates that produced significant metabolic signal over background, a t-test was performed individually on each of the A_750_-subtracted maximum absorbance results for each substrate from the 0 uM CPC control group. Using data collected from 3-4 days of experiments, an unpaired parametric two-tailed t-test with 90% confidence (p ≤ 0.1) was used to compare max signal from each substrate to the max signal from the no-substrate well. Substrates which reached statistical significance are shown in Table S1 and were selected for further analysis.

**Table S2. Substrate Slope Parameters for Biolog Electron Flow Rate Assay Analysis**

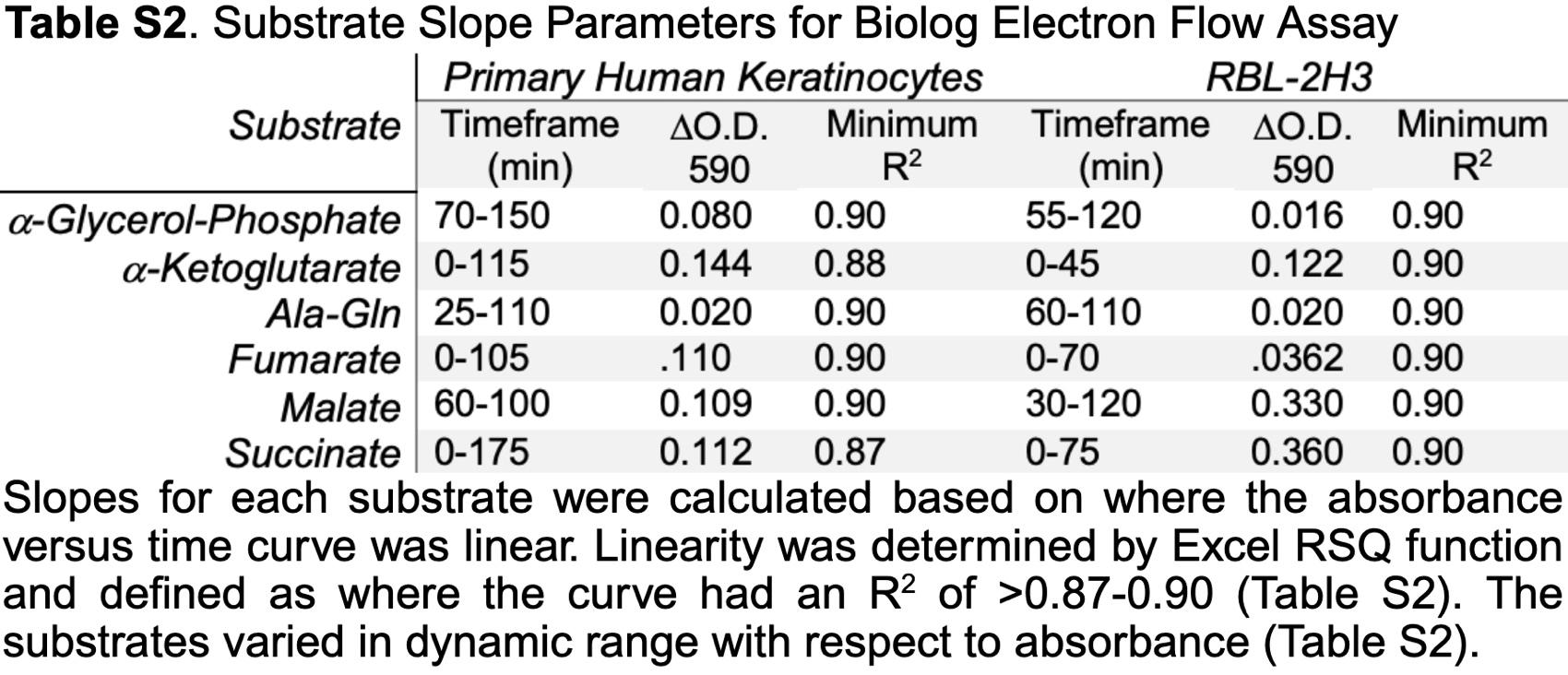

**Fig. S2. Effect of CPC on TCA cycle Enzyme Mitochondrial Malate Dehydrogenase (MDH2) Enzyme Activity**

**Method:** At substrate (malate) concentrations below 30 mM, MDH2 displays Michaelis-Menten kinetics (Raval et al., 1963; Telegdi et al., 1973). To assess the forward reaction of MDH2, concentrations of malate have been shown to be most effective at or below 8-10 mM (Heyde et al., 1968; Mueggler et al., 1978).

An MDH2 activity kit (Abcam) was employed. Pure human malate dehydrogenase (MDH2; CAS no. 9028-46-0; Novus) was prepared in kit-provided Tris buffer at 8 ug/mL. CPC was prepared as described in Materials and Methods at a 10X concentration. When added to the enzyme for pre-incubation, CPC is diluted to the final 1X concentration shown in Fig S2, and the MDH2 was 7.2 ug/mL. CPC is then preincubated for 60 min at 37 ºC/ 5 %CO_2_ with the enzyme. Kit reagents sodium malate, reagent dye (water-soluble tetrazolium redox dye), coupler (1-methoxy-5-methylphenazinium methyl sulfate), and NAD^+^ were diluted in Tris base buffer, pH 7.4 to compose the 2X activity solution, containing a consistent final 1:1:1:1 ratio of each reagent. Following the CPC pre-incubation, enzyme-CPC solutions were introduced to each substrate-containing activity solution in a half-area clear-bottom 96 well plate (VWR), resulting in a total volume of 100 uL per well (and MDH2 concentration 3.6 ug/mL) and promptly read in a plate reader at room temperature. The reporter dye is coupled to the production of NADH, resulting in a yellow color change measured by A_450_. The absorbance represents product formation and thus directly measures enzyme activity. Initial velocities were plotted as a function of malate concentration.

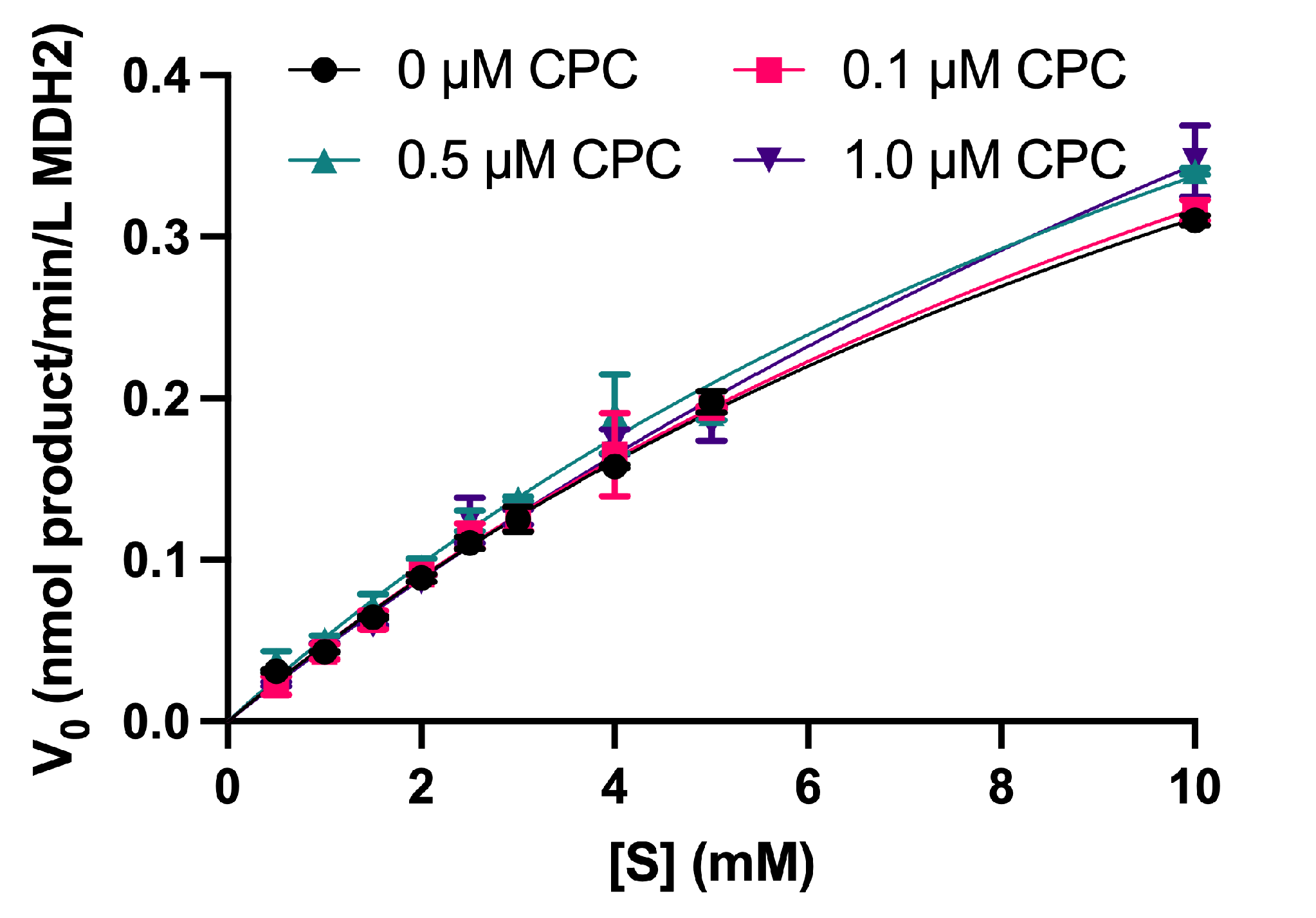

**Figure S2. CPC effects on enzyme activity of pure human malate dehydrogenase.** Purified MDH2 was assayed for activity with various substrate concentrations to create a Michaelis-Menten curve. Data shown are mean ± SEM. A one-way ANOVA with Tukey's post-hoc was used to determine significant differences. No significant differences were found.

**Results:** CPC (30 min, up to 1.0 uM) does not affect the enzymatic activity of MDH2.

**Conclusion:** The inhibition of electron flow from malate dehydrogenase through cytochrome c (Fig. 2, 3) is likely not due to direct inhibition of TCA enzyme MDH2. Interestingly, MDH2 is the second of the eight total TCA enzymes tested in this study. As with MDH2, succinate dehydrogenase was also unaffected by CPC (Complex II; Mitocheck). These findings suggest that the TCA cycle is not CPC’s direct target.

**Table S3. Readout Absorbances Used for Mitocheck Assays**

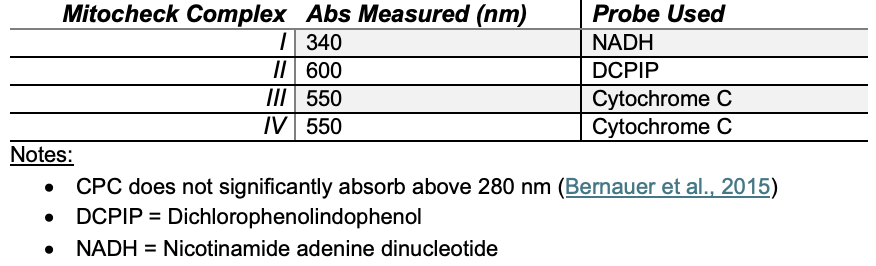

**Fig. S3. Justification for Mitocheck Complex I Conditions and Analysis Parameters: Time Range Selection, Absorbance Range Measured, and Vehicle Buffer** **Effects**

**Method:** Methods used to collect these data can be found in the Materials and Methods section of this manuscript. Note: for all Fig. 4 and Fig. S3 Mitocheck results, apart from Fig. S3B, DMSO at 0.05% was included in all assay solutions due to the amount of solvent required to fully dissolve the positive control rotenone.

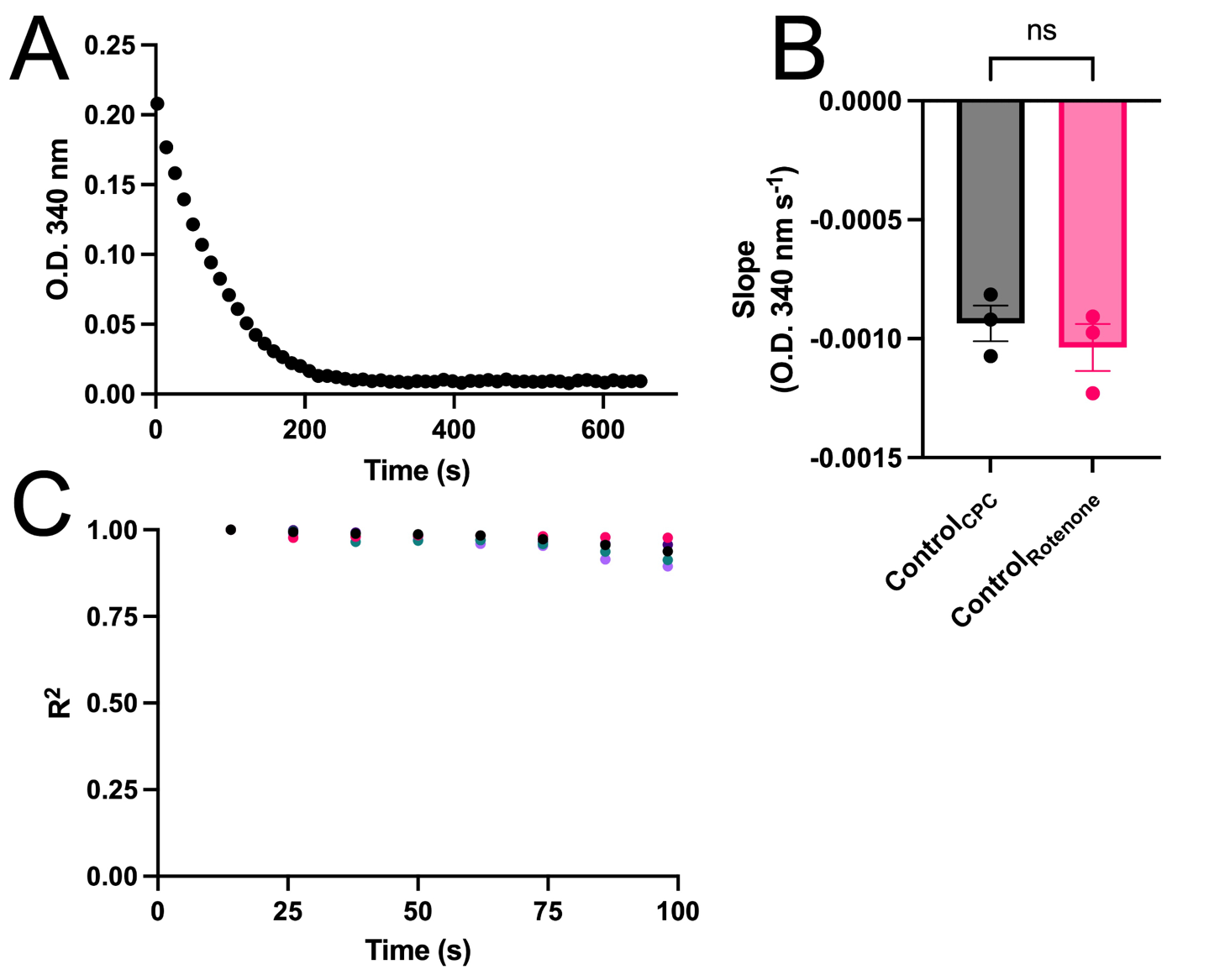

**Figure S3. Conditions and slope parameters for Mitocheck Complex I assay analysis.** **(A)** Representative graph of A_340_ for the control condition from one representative day. These A_340_ values were background subtracted, where the background was mitochondria in the Mitocheck buffer containing 0.05% DMSO and no additional NADH. **(B)** Slopes depicting the activities of Complex I in the absence of CPC, with either the 0.05% DMSO in the buffer (“Control_rotenone_”) or without DMSO (“Control_CPC_”). Data shown are mean ± SEM for 3 experiments; analysis was an unpaired T-test. **(C)** Assessment of linearity of readout response measuring complex I activity in untreated mitochondria. Linearity (R^2^) range of Complex I activity in the absence of CPC; linear time range determined by R^2^ > 0.90 threshold. Each plot represents one experimental day.

**Results:** Optimization of Mitocheck Complex I assay led to a robust range of A_340_ (Fig. S3A) in the linear range, R^2^ > 0.90 (Fig. S3C) of 98 sec. Additionally, there was no change in slope when comparing the control group in aqueous buffer compared to 0.05% DMSO buffer (Fig. S3B).

**Conclusion:** The time range and conditions used for Mitocheck Complex I data analysis represent data with a high linearity and a robust A_340_ range. Additionally, the presence of 0.05% DMSO in the assay buffer used did not affect Complex I activity, indicating that use of a standard buffer containing rotenone’s vehicle is justified for Complex I Mitocheck assays.

**Fig. S4. Justification for Mitocheck Complex II Conditions and Analysis Parameters: Time Range Selection, Absorbance Range Measured, and Vehicle Buffer** **Effects**

**Method:** Methods used to collect these data can be found in the Materials and Methods section of this manuscript. Note: for all Fig. 5 and Fig. S4 Mitocheck results, apart from Fig. S4B, DMSO at 0.14% was included in all assay solutions due to the amount of solvent required to fully dissolve the positive control TTFA.

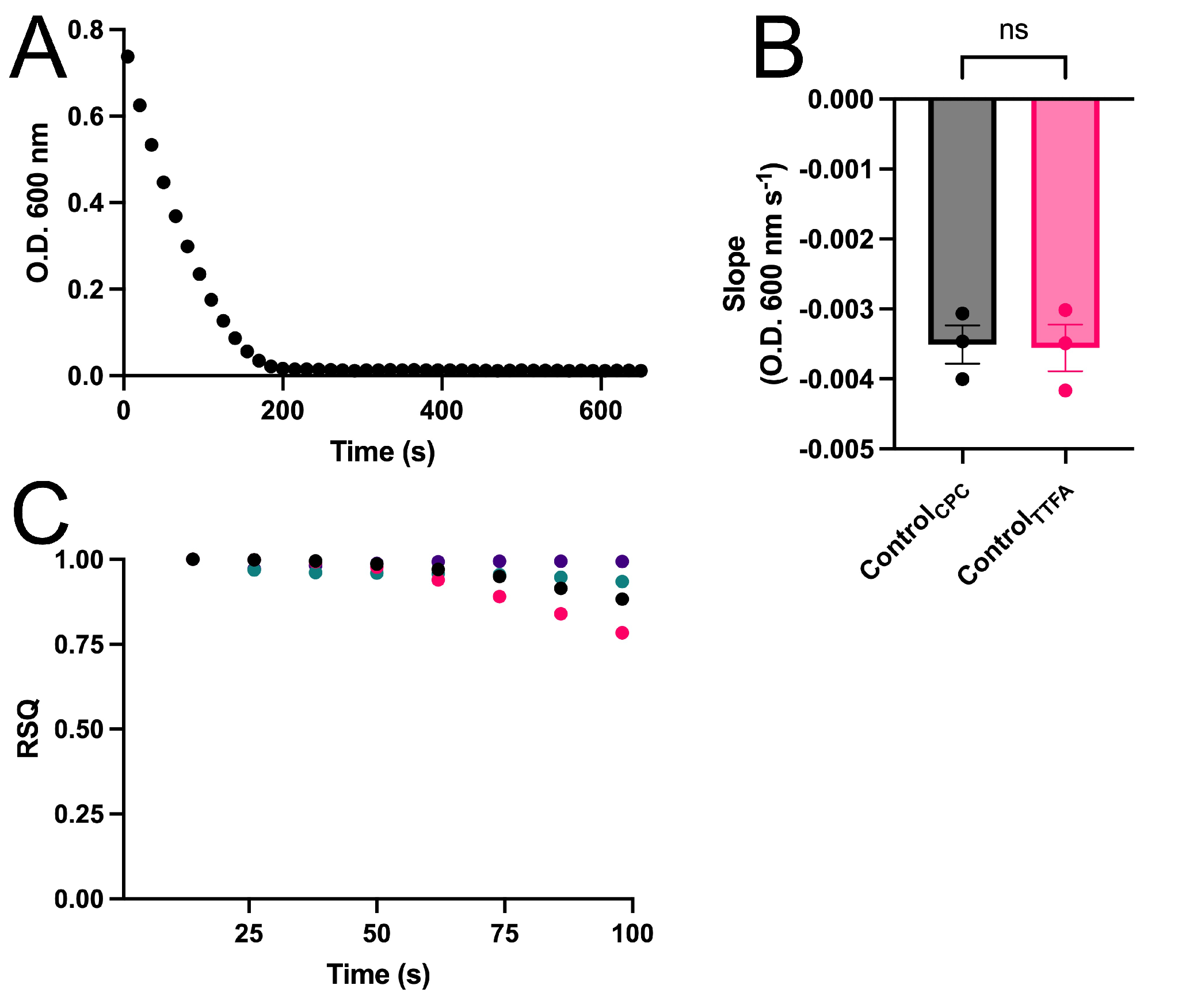

**Figure S4. Conditions and slope parameters for Mitocheck Complex II assay analysis.** **(A)** Representative graph of A_600_ for the control condition from one representative day. These A_600_ values were background subtracted, where the background was mitochondria in the Mitocheck buffer containing 0.14% DMSO and no additional succinate. **(B)** Slopes depicting the activities of Complex II in the absence of CPC, with either the 0.14% DMSO in the buffer (“Control_TTFA_”) or without DMSO (“Control_CPC_”). Data shown are mean ± SEM for 3 experiments; analysis was an unpaired T-test. **(C)** Assessment of linearity of readout response measuring complex II activity in untreated mitochondria. Linearity (RSQ) range of Complex II activity in the absence of CPC; linear time range determined by R^2^ > 0.95 threshold. Each plot represents one experimental day.

**Results:** Optimization of Mitocheck Complex II assay led to a robust range of A_600_ (Fig. S4A) in the linear range, R^2^ > 0.95 (Fig. S4C) of 86 sec. Additionally, there was no change in slope when comparing the control group in aqueous buffer compared to 0.14% DMSO buffer (Fig. S4B).

**Conclusion:** The time range and conditions used for Mitocheck Complex II data analysis represent data with a high linearity and a robust A_600_ range. Additionally, the presence of 0.14% DMSO in the assay buffer used did not affect Complex II activity, indicating that use of a standard buffer containing TTFA’s vehicle is justified for Complex II Mitocheck assays.

**Fig. S5. Justification for Mitocheck Complex III Conditions and Analysis Parameters: Time Range Selection, Absorbance Range Measured, and Vehicle Buffer** **Effects**

**Method:** Methods used to collect these data can be found in the Materials and Methods section of this manuscript. Note: for all Fig. 6 and Fig. S5 Mitocheck results, apart from Fig. S5B, DMSO at 0.28% was included in all assay solutions due to the amount of solvent required to fully dissolve the positive control antimycin a.

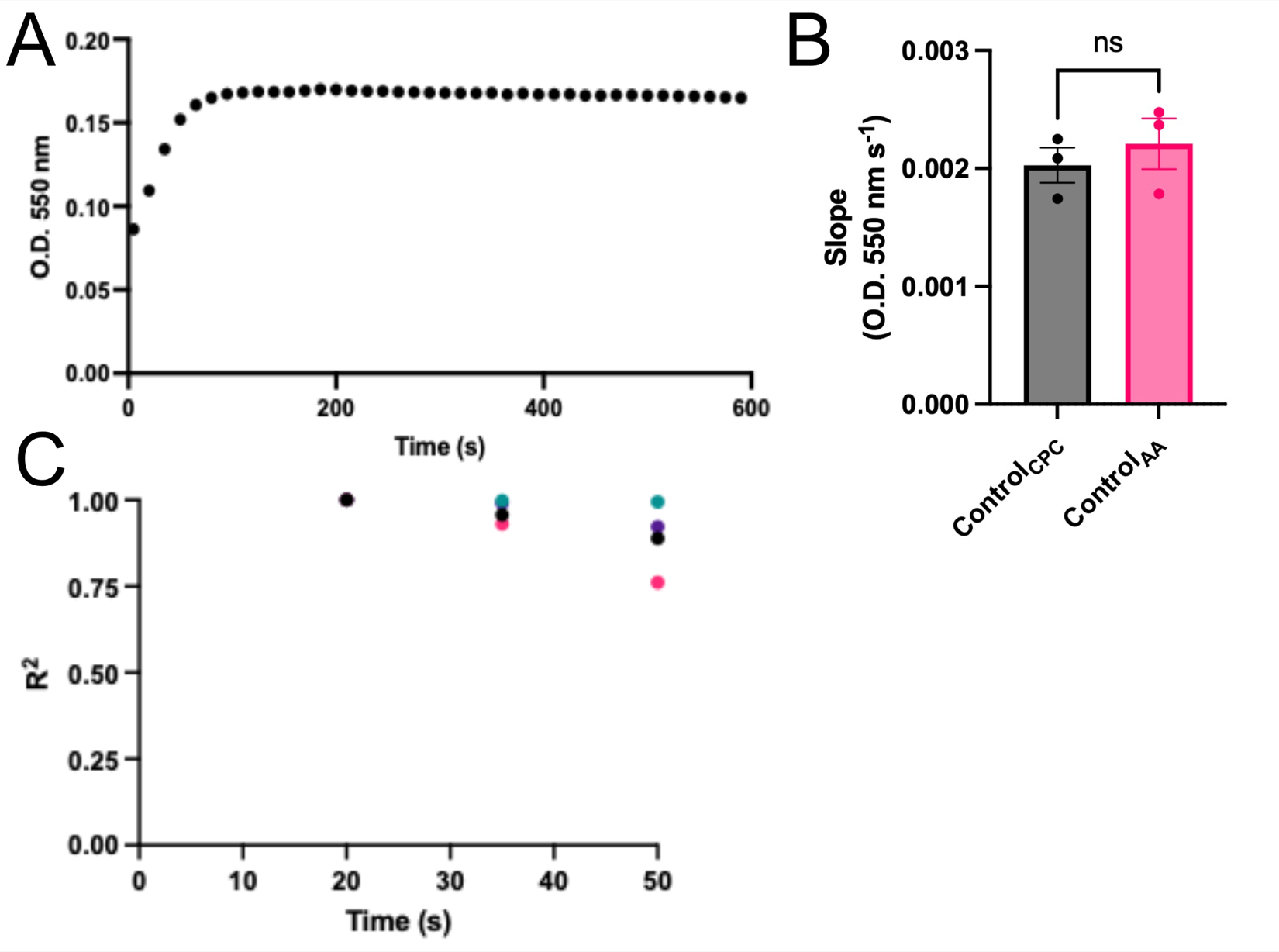

**Figure S5. Conditions and slope parameters for Mitocheck Complex III assay analysis.** **(A)** Representative graph of A_550_ for the control condition from one representative day. These A_550_ values were background subtracted, where the background was mitochondria in the Mitocheck buffer containing 0.28% DMSO and no additional succinate. **(B)** Slopes depicting the activities of Complex III in the absence of CPC, with either the 0.28% DMSO in the buffer (“Control_AA_”) or without DMSO (“Control_CPC_”). Data shown are mean ± SEM for 3 experiments; analysis was an unpaired T-test. (AA; antimycin a) **(C)** Assessment of linearity of readout response measuring complex III activity in untreated mitochondria. Linearity (R^2^) range of Complex III activity in the absence of CPC; linear time range determined by R^2^ > 0.80 threshold. Each plot represents one experimental day.

**Results:** Optimization of Mitocheck Complex III assay led to a robust range of A_550_ (Fig. S5A) in the linear range, R^2^ > 0.80 (Fig. S5C) of 50 sec. Additionally, there was no change in slope when comparing the control group in aqueous buffer compared to 0.28% DMSO buffer (Fig. S5B).

**Conclusion:** The time range and conditions used for Mitocheck Complex III data analysis represent data with a high linearity and a robust A_550_ range. Additionally, the presence of 0.28% DMSO in the assay buffer used did not affect Complex III activity, indicating that use of a standard buffer containing antimycin a’s vehicle is justified for Complex III Mitocheck assays.

**Fig. S6. Justification for Mitocheck Complex IV Conditions and Analysis Parameters: Time Range Selection, Absorbance Range Measured, and Vehicle Buffer** **Effects**

**Method:** Methods used to collect these data can be found in the Materials and Methods section of this manuscript. Note: for all Fig. 7 and Fig. S6 Mitocheck results, apart from Fig. S6B, 0.01 M NaOH was included in all assay solutions due to the amount of solvent required to fully dissolve the positive control KCN.

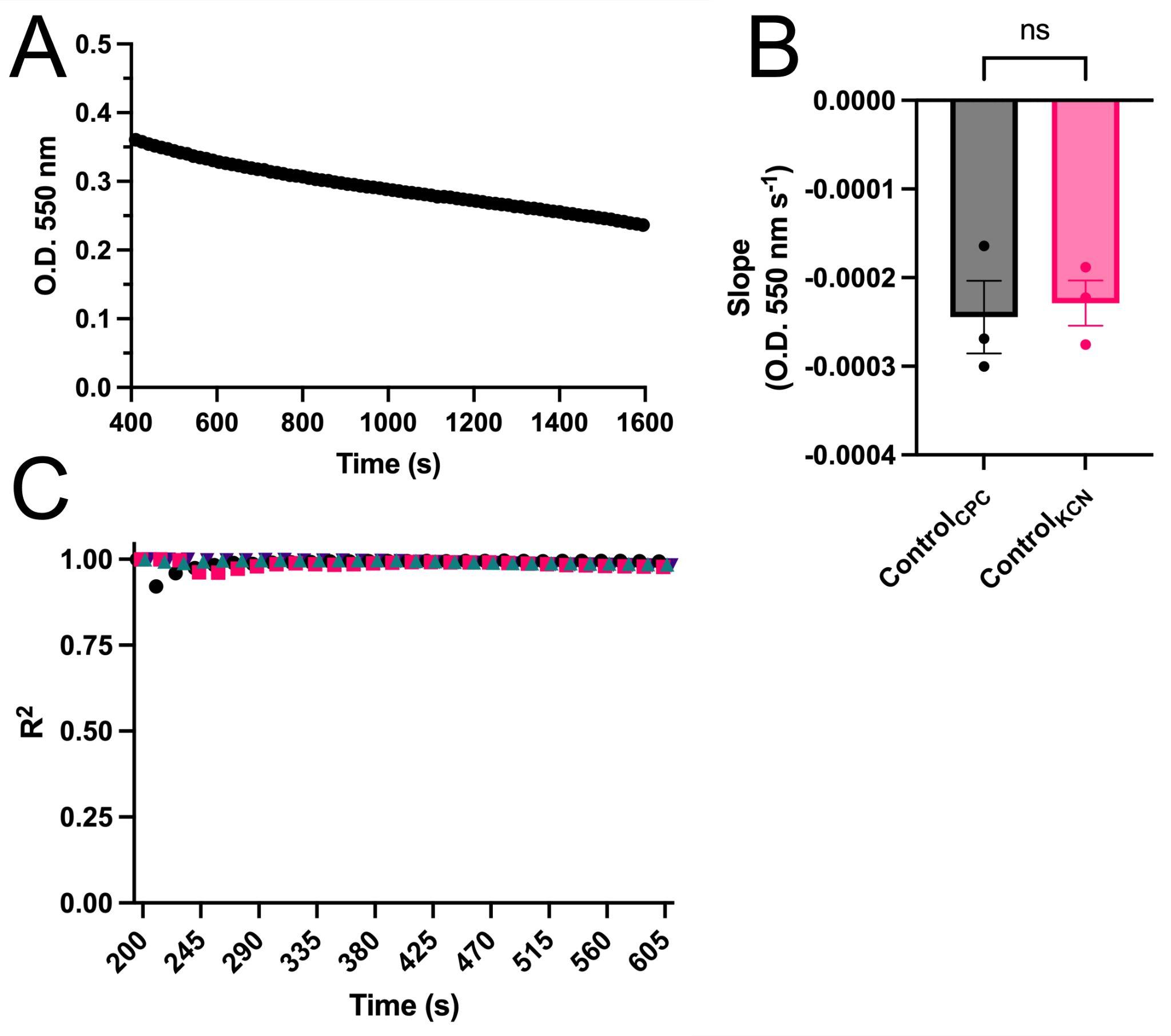

**Figure S6. Conditions and slope parameters for Mitocheck Complex IV assay analysis.** **(A)** Representative graph of A_550_ for the control condition from one representative day. These A_550_ values were background subtracted, where the background was mitochondria in the Mitocheck buffer containing 0.05 M NaOH and no additional cytochrome c. **(B)** Slopes depicting the activities of Complex IV in the absence of CPC, with either the 0.05 M NaOH buffer (“Control_KCN_”) or without NaOH (“Control_CPC_”). Data shown are mean ± SEM for 3 experiments; analysis was an unpaired T-test. **(C)** Assessment of linearity of readout response measuring Complex IV activity in untreated mitochondria in the Control_CPC_ buffer. Linearity (R^2^) range of Complex IV activity in the absence of CPC; linear time range determined by R^2^ > 0.9 threshold. Each plot represents one experimental day.

**Results:** Optimization of Mitocheck Complex IV assay led to a robust range of A_550_ (Fig. S6A) in the linear range, R^2^ > 0.9 (Fig. S6C) of 3-10 min. (When repeated in NaOH-containing Control_KCN_ buffer, the linear range was defined by R^2^ > 0.8, data not shown.) Additionally, there was no change in slope when comparing the control group in aqueous buffer compared to 0.05 M NaOH buffer (Fig. S6B).

**Conclusion:** The time range and conditions used for Mitocheck Complex IV data analysis represent data with a high linearity and a robust A_550_ range. Additionally, the presence of 0.05 M NaOH in the assay buffer used did not affect Complex IV activity, indicating that use of a standard buffer containing KCN’s vehicle is justified for Complex IV Mitocheck assays.

**Fig. S7. Total Cardiolipin Levels in RBL-2H3 Cells Fed Glucose-free Galactose-DMEM Media**

**Method:** In order to determine whether CPC interferes with levels of cardiolipin in RBL-2H3 mast cells fed galactose-DMEM media, the Abcam fluorescent cardiolipin assay was repeated (see “Materials and Methods”) using glucose-free galactose-DMEM media rather than the glucose-BT.

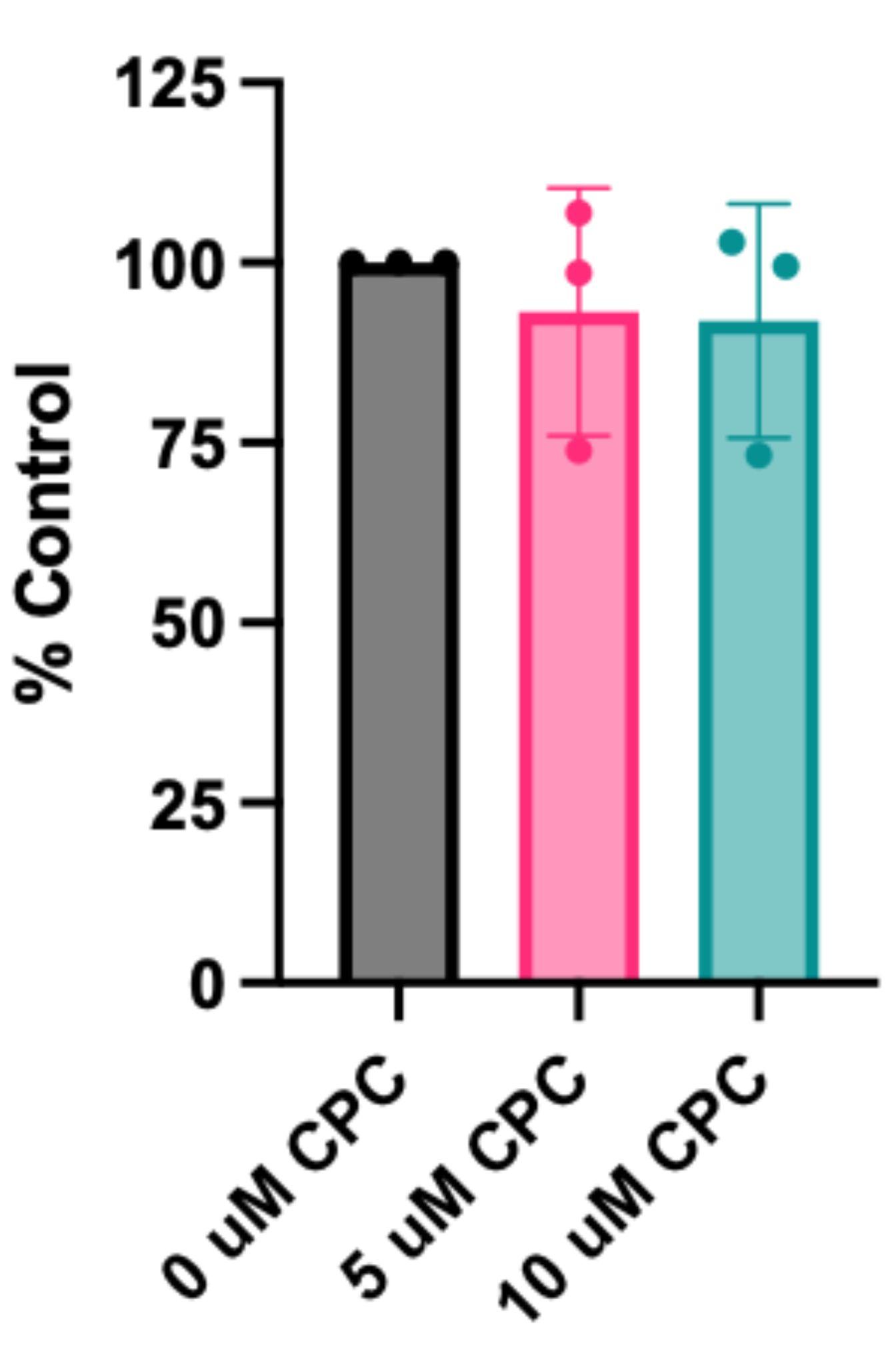

**Figure S7. Total cardiolipin levels in RBL-2H3 cells fed galactose-DMEM.** Intact RBL-2H3 cells were exposed to CPC for 1 h in DMEM-galactose, then lysed via sonication to determine cardiolipin levels via fluorescent dye. Lysates were analyzed via Bradford protein assay to standardize signals. Cardiolipin levels were normalized to the no-treatment control of each given day and are shown as mean ± SEM from 3 days of experiments (duplicates per experiment). Significance analysis was performed via one-way ANOVA with Dunnett's post-hoc test.

**Result:** CPC does not affect cardiolipin levels in RBL-2H3 cells under glucose-free conditions, using galactose-DMEM media.

**Conclusion**: Feeding cells galactose rather than glucose forces the cells to use their mitochondria for energy production (Lowry and Passonneau, 1969; Mulhausen and Mendicino, 1970; Rossignol et al., 2004). Therefore, even in cells undergoing significant mitochondrial function, total cellular levels of cardiolipin are not affected by CPC exposure.

Rossignol, R., Gilkerson, R., Aggeler, R., Yamagata, K., Remington, S.J., Capaldi, R.A.,

2004. Energy substrate modulates mitochondrial structure and oxidative capacity in

cancer cells. Cancer Res. 64, 985–993.

Telegdi, M., Wolfe, D.V., and Wolfe, R.G. (1973). Malate Dehydrogenase. J. Biol. Chem. *248*, 6484–6489. 10.1016/S0021-9258(19)43471-6.
